# Diverse plant RNAs coat Arabidopsis leaves and are distinct from apoplastic RNAs

**DOI:** 10.1101/2024.05.15.594325

**Authors:** Lucía Borniego, Meenu Singla-Rastogi, Patricia Baldrich, Megha Hastantram Sampangi-Ramaiah, Hana Zand Karimi, Madison McGregor, Blake C. Meyers, Roger W. Innes

## Abstract

Transgenic expression of a double-stranded RNA in plants can induce silencing of homologous mRNAs in fungal pathogens. Although such host-induced gene silencing is well-documented, the molecular mechanisms by which RNAs can move from the cytoplasm of plant cells across the plasma membrane of both the host cell and fungal cell are poorly understood. Indirect evidence suggests that this RNA transfer may occur at a very early stage of the infection process, prior to breach of the host cell wall, suggesting that silencing RNAs might be secreted onto leaf surfaces. To assess whether Arabidopsis plants possess a mechanism for secreting RNA onto leaf surfaces, we developed a protocol for isolating leaf surface RNA separately from intercellular (apoplastic) RNA. This protocol yielded abundant leaf surface RNA that displayed an RNA banding pattern distinct from apoplastic RNA, suggesting that it may be secreted directly from the leaf surface rather than exuded through stomata or hydathodes. Notably, this RNA was not associated with either extracellular vesicles or protein complexes; however, RNA species longer than 100 nucleotides could be pelleted by ultracentrifugation. Pelleting was inhibited by the divalent cation chelator EGTA, suggesting that these RNAs may form condensates on the leaf surface. These leaf surface RNAs are derived almost exclusively from Arabidopsis, but come from diverse genomic sources, including rRNA, tRNA, mRNA, intergenic RNA, microRNAs, and small interfering RNAs, with tRNAs especially enriched. We speculate that endogenous leaf surface RNA plays an important role in the assembly of distinct microbial communities on leaf surfaces.

**Significance Statement:** Plant leaves are colonized by a complex community of microbes that is shaped by host genetics. Although secreted metabolites are thought to mediate this effect, we investigated whether plants might also secrete RNA that could potentially structure microbial communities via cross-kingdom RNA interference. Here we report that Arabidopsis leaves are covered with diverse RNAs of plant origin, including abundant tRNAs and tRNA fragments. This leaf surface RNA is not associated with extracellular vesicles or protein complexes; however, it is less degraded than RNA found inside the extracellular spaces of leaves, suggesting that leaf surface RNA is secreted directly rather than exuded through stomata or hydathodes. We propose that this RNA plays a direct role in shaping the leaf microbiome.

## Introduction

Secretion of RNA into the extracellular environment is a well-conserved phenomenon as it is known to occur in all life forms (1–6). Recent studies in both mammalian and plant systems have revealed that the extracellular RNA (exRNA) pool is highly diverse, with the majority of exRNA associated with RNA-binding proteins outside of extracellular vesicles (EVs) (7–9). However, there are only a few reports in plants wherein exRNAs other than those encapsulated inside EVs have been described (10, 11).

Most studies on plant exRNA have focused on small non-coding RNAs, including microRNAs (miRNAs) and small interfering RNAs (siRNAs), due to their roles in RNA interference (RNAi)-mediated gene silencing. The RNAi pathway is highly conserved and is triggered by dsRNA molecules that are recognized and processed into siRNAs by Dicer-like proteins (DCLs). The resulting siRNAs bind to Argonaute proteins (AGOs) to form RNA-induced silencing complexes (RISC), and subsequently, the siRNAs guide the RISC to target mRNA transcripts for degradation (12–15).

Two technologies that exploit RNAi to protect plants against invading phytopathogens have been successfully implemented. Host-induced gene silencing (HIGS) involves the incorporation of a transgene expressing a double-stranded RNA (dsRNA) in a plant that targets an essential pathogen gene (11). Spray-induced gene silencing (SIGS), in contrast, involves exogenous application of dsRNAs or siRNAs onto foliar surfaces. Both approaches have been shown to induce sequence-specific gene silencing in microbial pathogens, insects, and nematodes (16, 17). Notably, dsRNA sprayed onto barley leaves leads to the inhibition of fungal growth in non-sprayed distal tissues (16). Although the accumulation of unprocessed dsRNA was confirmed in distal tissues, the corresponding siRNAs were missing, suggesting that dsRNAs can be translocated within or on a plant leaf without being processed by DCLs, possibly in the apoplast.

One potential mechanism for translocation of silencing RNAs (miRNAs, siRNA, and dsRNAs) is via packaging inside extracellular vesicles (EVs). Two studies have reported the uptake of plant EVs carrying small RNAs by fungal cells, both *in vitro* and *in vivo* (18, 19); however, strong evidence supporting this mechanism of RNA uptake is still lacking. Although selective packaging inside plant EVs is the most widely studied trafficking system for silencing RNAs (10, 17–21), how RNAs are packaged inside EVs and how they are then taken up by pathogens is poorly understood. Additionally, if EVs are taken up via endocytosis, how silencing RNAs can then escape the endosomes to engage the pathogen RNAi machinery is unknown.

In a recent report, Schlemmer et al. (22) isolated plant extracellular vesicles (EVs) from both dsRNA-expressing transgenic Arabidopsis plants and dsRNA-sprayed barley leaves to assess their effect on fungal growth. Surprisingly, both sources of plant EVs carried very small amounts of dsRNA-derived siRNAs and did not affect fungal growth, indicating a minor role of EVs in both HIGS and SIGS (22). There are no reports thus far confirming the presence of dsRNA inside EVs. In another study, the exogenous application of sRNAs or dsRNAs targeting *Botrytis* DCL1 and DCL2 genes onto the surface of fruits, vegetables, and flowers was shown to significantly inhibit grey mold disease, suggesting that the pathogen is fully capable of taking up naked RNA (17).

Of particular note, HIGS-mediated resistance to a fungal pathogen appears to involve uptake of silencing RNAs by the fungal cells prior to penetrating plant cell walls (23). For instance, transgenic rice lines expressing silencing RNAs targeting the *MoAP1* gene of the fungal pathogen *Magnaporthe oryzae* displayed enhanced resistance to infection by *M. oryzae*. Although the fungal conidia germinated and formed appressoria, few of these formed infection hyphae, indicating that the fungus was blocked prior to its penetration into the plant cell wall. Similar observations were made for other fungal pathogens, including *Puccinia triticina* (24) and *Puccinia striiformis* f. sp. tritici (25, 26), in which genes highly expressed in early stages of infection were targeted by HIGS, thereby resulting in robust resistance. This suggests that the silencing RNAs are taken up by the fungi before haustoria formation, possibly from the leaf surface during the growth of the germ tube. In support of this hypothesis, in the above SIGS study conducted in barley, the dsRNA-treated leaves restricted the fungal mycelia to the inoculation sites, leaving the surrounding leaf tissue free of infection hyphae. More importantly, fungal germination was found to be strongly impaired (16). Together, these observations led us to hypothesize that silencing RNAs produced by plants may be deposited onto the plant leaf surface, an extracellular fraction that has been overlooked in previous RNAi studies.

Here we report that Arabidopsis leaf surfaces are coated with abundant RNA. Leaf surface RNA differs from apoplastic and cellular RNA both in composition and size. Furthermore, we found that the leaf surface RNA is not protected from endonuclease degradation either by EVs or RNA-binding proteins. In both the apoplast and leaf surface, we found tRNAs to be the most abundant, but leaf surface tRNAs were mostly intact, whereas apoplastic tRNAs were usually processed into tRNA halves (mainly produced by cleavage in the anticodon loop) and tRNA fragments (tRFs; produced by cleavage in the D loop). Such tRNA-derived molecules are now known to play gene regulatory roles in the context of plant-microbe interactions, which is independent of canonical tRNA function in translation (27, 28). We thus speculate that these extracellular tRNA-derived molecules may play a significant role in plant-microbe interactions. Additionally, we noted the enrichment of specific classes of miRNAs and siRNAs in the two extracellular fractions, which might also play a role in structuring the leaf microbiome and/or in mediating immune responses. Lastly, we found that leaf surface RNA forms cation-dependent condensates, which may contribute to the stability of these naked RNAs.

## Results

### Arabidopsis Secretes RNA onto the Leaf Surface

In our recent study, we demonstrated the presence of diverse species of RNA in the apoplast of Arabidopsis rosettes (11). To determine whether Arabidopsis plants also secrete RNA onto their leaf surface, we developed a method to collect both leaf surface wash (LSW) and apoplastic wash fluid (AWF) from the same set of plants. To collect LSW, we detached whole rosettes and sprayed both adaxial and abaxial leaf surfaces with vesicle isolation buffer (VIB) supplemented with 0.001% Silwet, followed by spinning at a very low speed (100 *g*) to avoid any contamination with apoplastic fluid or cellular damage. To isolate AWF, we vacuum infiltrated the same set of plants with VIB followed by centrifugation at 600 *g* (*SI Appendix*, Fig. S1). To avoid microbial contamination, these two extracellular fractions were filtered through 0.2 µm filters before processing. The collected fractions were clear, indicating that no cellular damage was caused during this isolation process. To confirm this, we stained the leaves with trypan blue dye, which stains ruptured cells. No ruptured cells were observed (*SI Appendix*, Fig. S2). We then isolated RNA directly from the filtered LSW and AWF, without conducting any ultracentrifugation steps. Analysis of the purified RNA using denaturing RNA gel analysis revealed the presence of diverse species of long and small RNAs in both LSW and AWF (Fig. 1 *A* and *B*). Notably, RNA amounts isolated from leaf surfaces were equivalent to those isolated from apoplastic fluids (normalized per plant fresh weight) (Fig. 1 *C* and *D* and *SI Appendix*, Fig. S3). However, the RNA size profiles of the LSW and AWF fractions were substantially different from each other, and these two profiles differed from those observed for total cell lysate (CL) RNA, indicating that the two extracellular fractions were not contaminated with cellular RNA. Overall, apoplastic RNA was the most diverse fraction in terms of its size distribution. Although both extracellular fractions were enriched in RNAs smaller than 80 nt compared to total CL, species smaller than 60 nt were especially abundant in AWF. Of note, there was a strong accumulation of RNAs ranging from 30 to 35 nt in length in AWF, which were not as conspicuous in LSW. We observed a similar trend for tiny RNAs (<18 nt) (Fig. 1*A*). The differences in the banding pattern between LSW and AWF were also quite prominent in RNAs longer than 150 nt (Fig. 1*B*). The overall banding pattern of long RNAs in LSW was more similar to that of CL than that of AWF. For instance, RNAs longer than 1000 nt were present in both LSW and CL, while they were absent in the AWF samples. These differences in the banding pattern between LSW and AWF indicate that RNA on the leaf surface is unlikely to be derived from the apoplast, and thus is unlikely to be secreted through stomata or hydathodes.

**Fig. 1.**
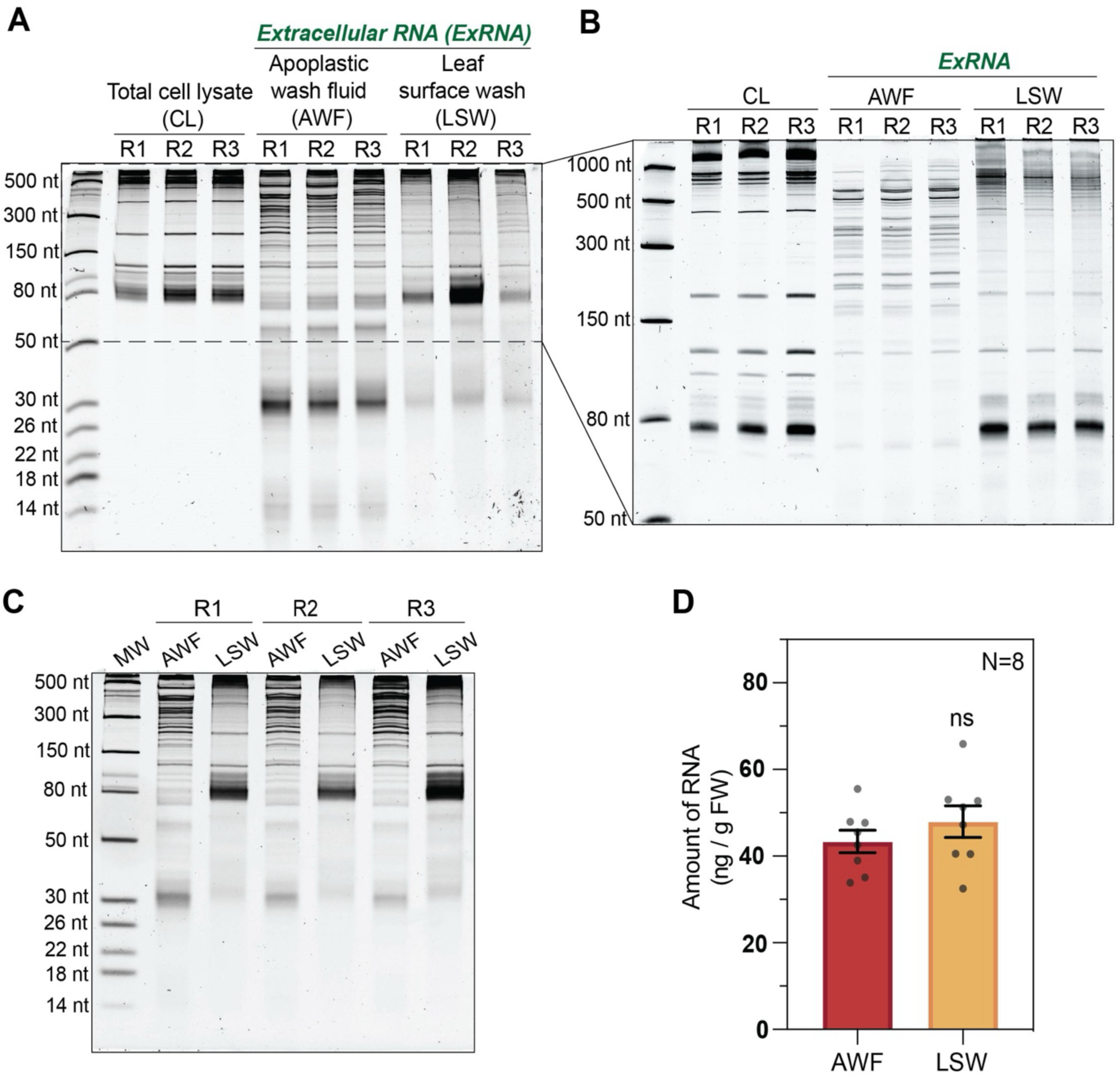
Plant leaves are coated with RNA. (*A*) Five-to-six-week-old Arabidopsis plants were used to collect leaf tissue (CL), apoplastic wash fluid (AWF), and leaf surface wash (LSW) from three biological replicates. RNA was isolated using TRIzol, and 100 ng of RNA from the cellular and extracellular fractions (AWF and LSW) was separated on a 15% denaturing polyacrylamide gel and stained with SYBR GOLD nucleic acid stain to visualize and compare the RNA-banding pattern between CL, AWF, and LSW. RNA size standards are shown in the leftmost lane. (*B*) CL, AWF, and LSW RNA (100 ng) was separated on a 10% denaturing polyacrylamide gel and stained with SYBR GOLD nucleic acid stain to better resolve the RNA between 50–1000 nt. (*C*) To estimate the concentration of RNA in AWF and LSW isolated per gram fresh weight (ng/g FW) of leaf tissue, RNA was separated on a 15% denaturing polyacrylamide gel and densitometry analysis was performed using ImageJ software. (*D*) Bar graph represents the normalized amount of RNA isolated from eight independent replicates (N=8; R4-R8 shown in *SI Appendix,* Fig. S2) of AWF and LSW plotted with mean and standard error. Student’s t-test was performed to determine statistical differences in RNA concentration of AWF and LSW. ns: not significant.

### Unlike Apoplastic RNA, Leaf Surface RNAs Are Not Associated with Proteins and Can Be Fully Degraded by Endoribonucleases

We have previously reported that RNA purified from P40 pellets isolated from AWF (obtained by centrifuging AWF at 40,000 *g*) is partially resistant to RNase A treatment, most likely due to its association with proteins, but not due to its packaging inside EVs (11). To analyze the role of proteins in protecting exRNAs from ribonuclease digestion, we collected LSW and AWF from Arabidopsis rosettes as described above and performed an RNase A protection assay followed by RNA isolation using TRIzol (Fig. 2*A* and *SI Appendix*, Fig. S4). Denaturing polyacrylamide gel electrophoresis of these RNAs revealed that approximately 85–90% of the RNA in AWF was readily digested after treatment with 9 µg/mL of RNase A at room temperature (RT) for 1 h, while all the RNA was digested when AWF was pretreated with trypsin before RNase A treatment. These results are consistent with our previous reports using P40 pellets and indicate that proteins do play a role in preventing apoplastic RNA from being digested by endoribonucleases (11). Although AWF RNA was partly protected from digestion by RNase A, there was almost complete digestion of longer RNAs, and we noticed a change in the size distribution of RNAs smaller than 150 nt. Such alteration is indicative of partial digestion of RNAs, revealing an incomplete protection of RNA molecules. To analyze the degree of RNA protection by proteins in the AWF, we treated AWF with different concentrations of RNase A (*SI Appendix*, Fig. S5). We observed a greater degradation of RNA with an increase in the concentration of RNase A (from 5– 20 µg/mL), but the overall banding pattern remained the same.

**Fig. 2.**
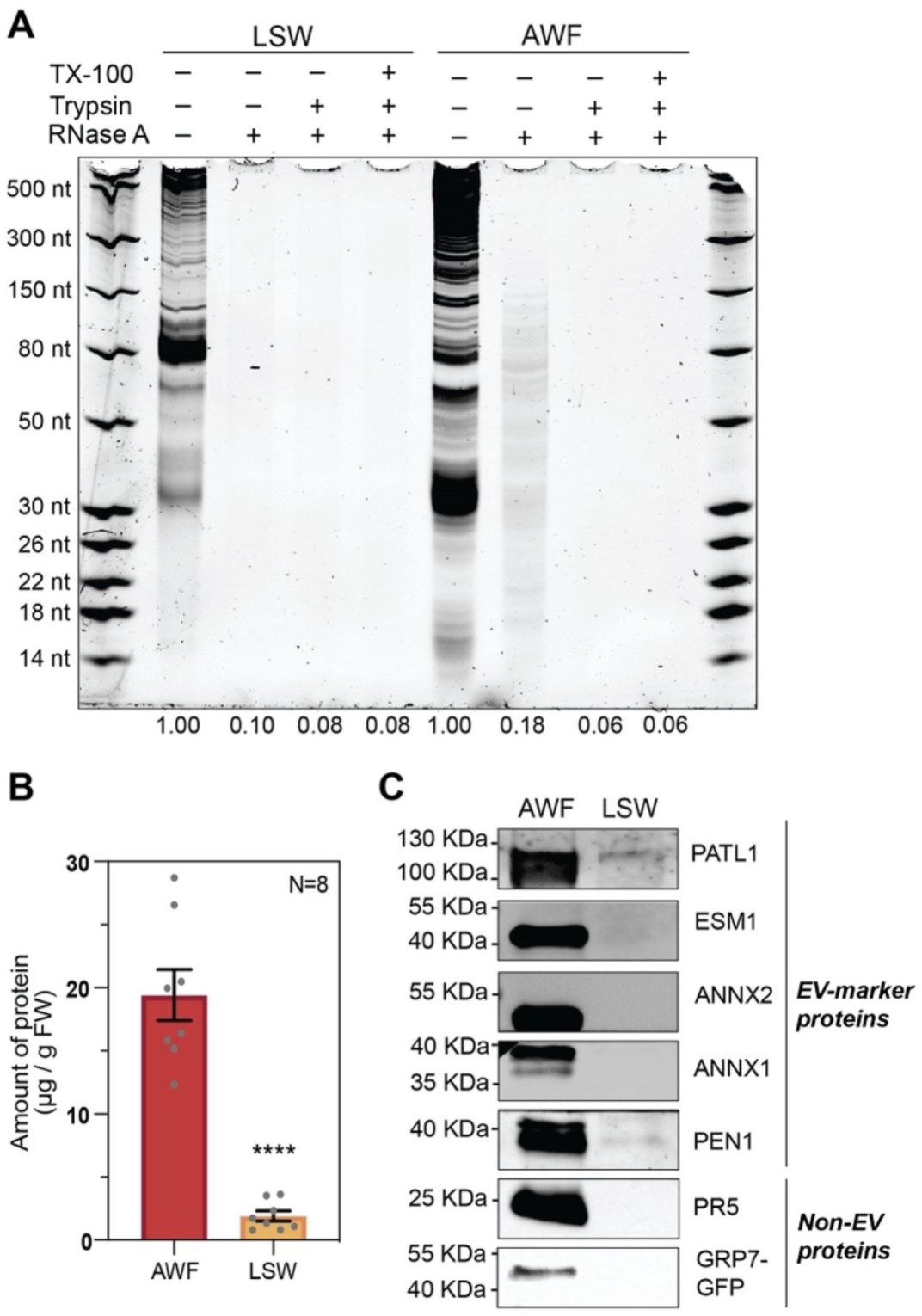
Unlike AWF, LSW fractions are depleted in proteins, leaving their RNAs unprotected from endoribonuclease digestion. (*A*) Ribonuclease protection assay of AWF and LSW. AWF and LSW samples were treated with RNase A, or trypsin followed by RNase A, or TX-100 followed by trypsin followed by RNase A. The negative control was mock-treated and kept on ice. RNAs were extracted using TRIzol, separated in a 15% denaturing polyacrylamide gel, and stained with SYBR Gold nucleic acid stain. RNA abundance in each gel lane was estimated by densitometry and expressed relative to the negative control (no RNase). This experiment was repeated three times on different days using three biological replicates (other replicates are shown in *SI Appendix,* Fig. S3). (*B*) Amount of protein in AWF and LSW expressed as µg of protein per gram of leaf fresh weight. The amount of protein in each sample was estimated by densitometry using the silver-stained gels presented in *SI Appendix,* Fig. S5. The graph shows the mean±SEM. Values from 8 biological replicates were normally distributed and significant differences were calculated using Welch’s t-test: ****, P <0.0001. (*C*) Immunodetection of EV-marker proteins (PEN1, PATL1, ESM1, ANNX1, ANNX2) and RNA-binding proteins (GRP7, PR5, ANNX1, ANNX2) in AWF and LSW fractions. The amount of protein loaded per fraction was normalized by leaf FW.

To assess whether LSW RNA is also partially protected by proteins, we performed a similar RNase A protection assay using 9 µg/mL of RNase A. To our surprise, RNase A alone was sufficient to fully digest all RNA in the LSW (Fig. 2*A* and *SI Appendix*, Fig. S4), indicating that RNAs on the leaf surface are not protected by either proteins or EVs.

### LSW Contains Few Proteins, Including EV-Associated Proteins

The observation that RNAs on the leaf surface are not protected from endoribonuclease digestion led us to test the abundance of EV-marker and RNA-binding proteins associated with LSW RNA. We loaded an equivalent volume of AWF and LSW (normalized per plant fresh weight) from eight different replicates and resolved proteins on an SDS-PAGE gel followed by silver staining (*SI Appendix*, Fig. S6*A*). Densitometric analysis revealed that there was substantially less protein in LSW compared to AWF (Fig. 2*B* and *SI Appendix*, Fig. S6*A*). We performed immunoblot analysis to specifically assay the EV-marker proteins PENETRATION 1 (PEN1), PATELLIN 1 (PATL1), and EPITHIOSPECIFIER MODIFIER 1 (ESM1) (21) in AWF and LSW (Fig. 2*C* and *SI Appendix*, Fig. S6*B*). We also assayed RNA-binding proteins, including ANNEXIN 1 and 2 (ANN1 and ANN2) that are known to be secreted by exosome-like EVs and bind to sRNAs non-specifically (20). We also probed for the non-EV secreted RNA binding proteins PATHOGENESIS-RELATED GENE 5 (PR5) and GLYCINE-RICH RNA-BINDING PROTEIN 7 (GRP7) (29–31). Surprisingly, LSW contained a very small amount of these proteins relative to AWF, which is consistent with the overall protein accumulation pattern observed in LSW (Fig. 2 *B* and *C* and *SI Appendix*, Fig. S6). These results confirm the absence or very low abundance of RNA-binding or EV-marker proteins in LSW. This lack of proteins on the leaf surface suggests that LSW may also lack RNases, which could account for the lower level of processing observed in RNA isolated from the leaf surface compared to that isolated from the apoplast (Fig. 1 *A* and *B*).

### AWF and LSW Exhibit a Distinct Small RNA Composition Compared to Total CL

To compare the content of leaf surface RNA to apoplastic and cellular RNA, we performed sRNA sequence analysis. We sequenced sRNA from three biological replicates of each fraction, including total CL, AWF, and LSW (a total of nine libraries). We observed that the distribution of read lengths was consistent between the three replicates, but there was a significant difference between the size distributions of total CL, AWF, and LSW RNA (Fig. 3*A*). Total CL RNA displayed two predominant peaks at 21 and 24 nt, mostly corresponding to miRNAs and heterochromatic siRNAs (*SI Appendix*, Fig. S7). However, AWF and LSW RNA exhibited different size distribution patterns with peaks at 16 and 31 nt, and 16, 20, and 32 nt, respectively (Fig. 3*A*). To understand the nature of these AWF and LSW sRNAs, we analyzed their genomic origin. We observed that most of the sRNAs in the extracellular fractions originated from rRNAs, cDNA, tRNAs, and products that were dependent on RNA polymerase IV (Pol IV) (Fig. 3*B* and *SI Appendix*, Fig. S9). We also observed that the AWF and LSW RNAs exhibited statistically significant enrichment in sRNAs derived from tRNAs when compared to total CL and statistically significant depletion in miRNAs, tasiRNAs, and rRNA-derived fragments (*SI Appendix*, Fig. S9). These accumulation patterns support our previous findings that exRNA is enriched in tRNA-derived fragments (10, 11). These observations are similar to those from mammalian systems, which have found that the majority of the exRNAome is comprised of fragments derived from rRNAs and tRNAs and are secreted into the extracellular milieu independent of EVs (8).

**Fig. 3.**
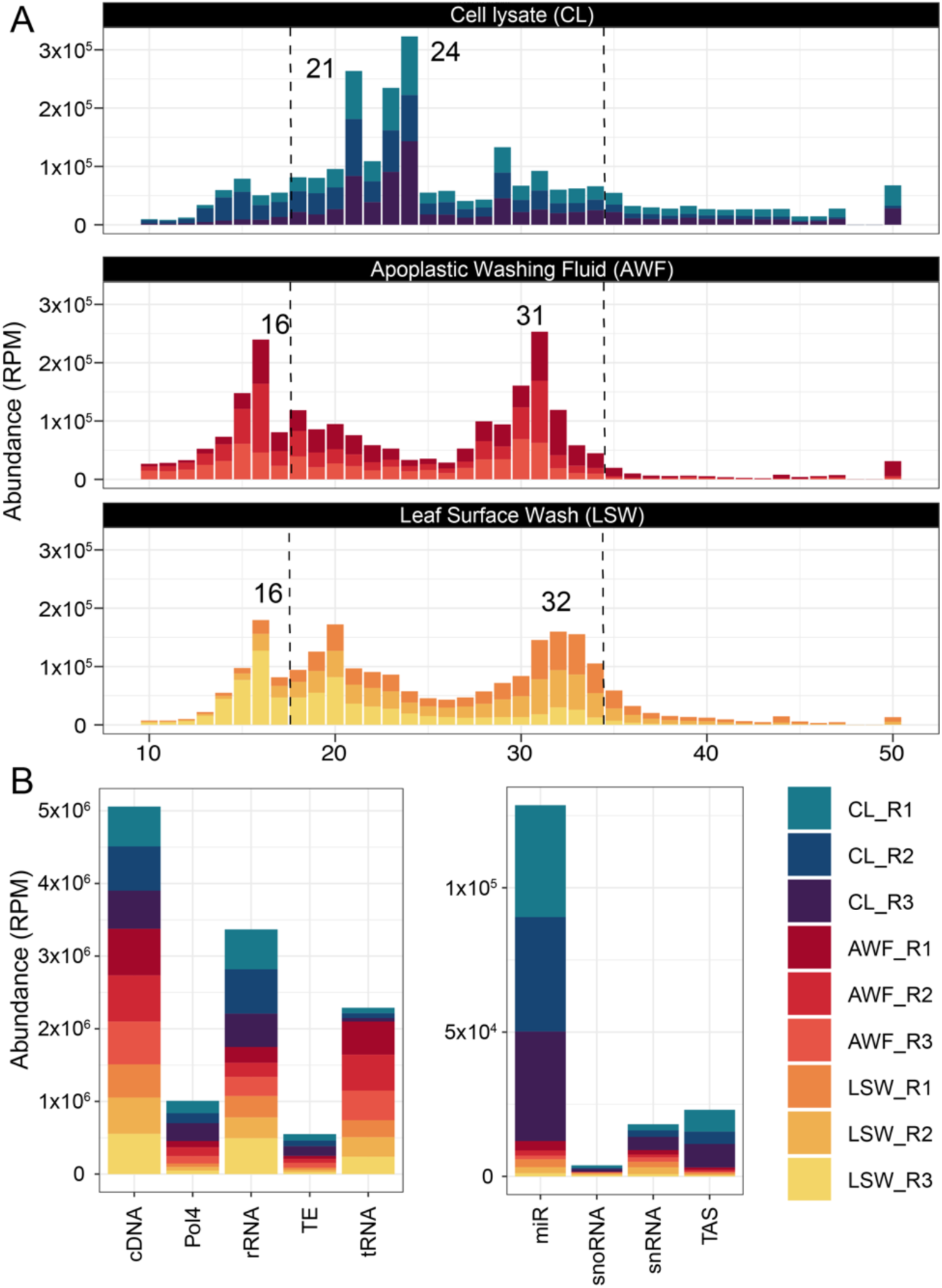
exRNA exhibits a distinct sRNA composition compared to cell lysate. (*A*) sRNA size distribution of reads mapping to the Arabidopsis genome (TAIR version 10). The abundance of each size class was calculated for each sample independently, and normalized to the total number of library reads. The *x*-axis indicates the sRNA size, from 10 to 50 nt long, and the *y*-axis indicates its abundance in reads per million (RPM) reads. The dotted lines delimit the consensus size ranges of small RNAs, from 18 to 34 nt (*B*) Genomic origin of small RNA reads based on the categories established in the TAIR 10 genome version. RNAs that mapped to the genome were categorized by origin and plotted by relative abundance in RPM. The x-axis represents the feature of interest, the left panel for features with higher abundance, and right panel for features with lower abundance. cDNA, complementary DNA; Pol4, Polymerase IV-dependent products; rRNA, ribosomal RNAs; TE, Transposable elements; tRNA, transfer RNAs; miR, microRNAs; snoRNA, small nucleolar RNA; snRNA, small nuclear RNAs; TAS, trans-acting siRNA. The three colored shades represent three independent biological replicates, with each replicate derived from 18 Arabidopsis plants.

### Specific miRNAs and tasiRNAs Differentially Accumulate in LSW and AWF

Even if miRNAs are generally underrepresented in the AWF and LSW fractions compared to total CL RNA, we wished to learn whether there were specific miRNAs that differentially accumulated in any of these three fractions. We observed that the size distribution of reads mapping to miRNAs was different in each fraction, shifting from mainly 21 nt in CL to 18 nt in AWF, and 20 nt in LSW (*SI Appendix*, Fig. S7*A*). Since the size of the miRNA influences AGO loading and its subsequent activity, we performed a differential expression analysis considering only reads that mapped to known miRNAs with lengths 20, 21, or 22 nt long. We observed that out of the 361 miRNAs detected, 128 differentially accumulated in at least one of the three comparisons (*SI Appendix*, Fig. S8). Based on their significance and accumulation level, these miRNAs were clustered into five groups. Cluster 1 contains 58 miRNAs that exhibit a higher accumulation in AWF and LSW when compared to CL. Out of these, 21 exhibit a higher accumulation in LSW compared to AWF. Cluster 2 contains 19 miRNAs that are significantly enriched in LSW and not in AWF when compared to CL and enriched in LSW when compared to AWF. Cluster 3 contains 23 miRNAs enriched in LSW compared to CL, depleted in AWF compared to CL, and enriched in LSW when compared to AWF. In other words, the 42 miRNAs included in Clusters 2 and 3 are enriched in LSW compared to the other two fractions. Cluster 4 contains 10 miRNAs that are depleted in both AWF and LSW compared to CL and enriched in LSW compared to AWF. Cluster 5 contains 18 miRNAs that are depleted in LSW when compared to both CL and AWF. Taken together, the miRNAs in clusters 4 and 5 exhibit a higher accumulation inside of cells than in the extracellular space.

When calculating miRNA differential accumulation, we considered relative proportions and not absolute accumulations; thus, a miRNA can be enriched in AWF and LSW compared to total CL, but have fewer absolute counts in AWF and LSW than in total CL. However, we also observed that 19 miRNAs had a higher number of reads in AWF and/or LSW, compared to total CL (Fig. 4). Out of these, eight miRNAs, belonging to three families, miR8167, miR5653, and miR5659, had a higher abundance in AWF than in LSW and CL (Fig. 4*A*). The other eleven miRNAs had a higher abundance in LSW than in the other two fractions. These miRNAs belong to six distinct, but highly conserved miRNA families, including miR156, miR169, miR172, miR5014, miR773, and miR829 (Fig. 4*B*). Of particular note, plants overexpressing miR156 have been found to secrete this miRNA into the growth medium, which is then taken up by wild-type plants in co-culture, resulting in downregulation of target genes (32). Since miRNAs can be functionally exchanged between neighboring plants via the environment, they are potential signaling agents. A plethora of putative functions could be ascribed to the extracellular miRNAs based on the current literature; however, further analysis is required to assign their biological functions.

**Fig. 4.**
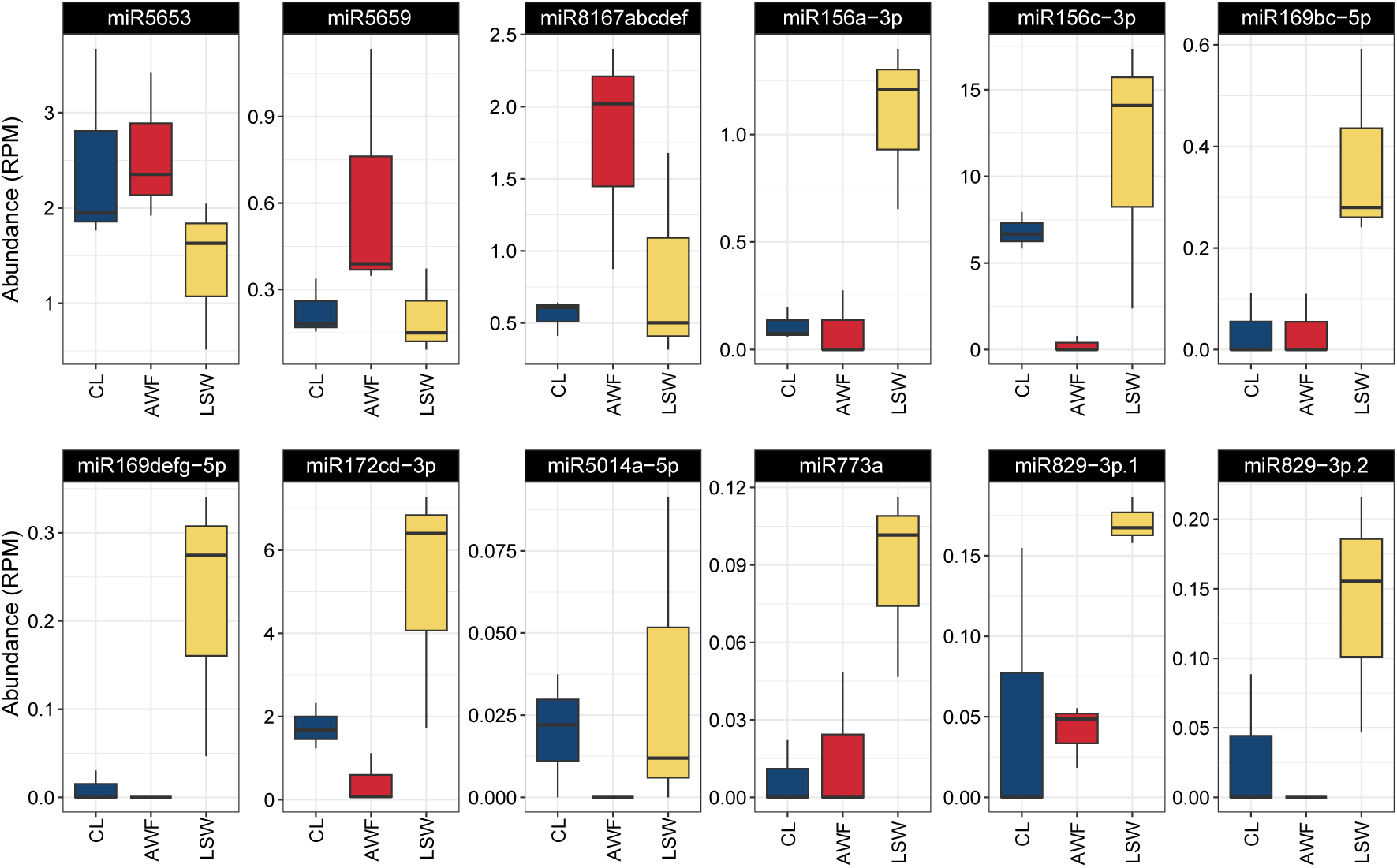
Specific miRNAs are more abundant in AWF or LSW compared to cell lysate. The first three box plots show miRNAs that are more abundant in AWF while the remaining show miRNAs that are more abundant in LSW. The x-axis represents the three distinct fractions, and the y-axis represents the absolute abundance in reads per million library reads (RPM). Each boxplot represents three biological replicates, where the box comprises data points between quartiles 1 and 3, and the whiskers show the sample variability outside of these quartiles. miRNA names with multiple letters (e.g., miR8167) indicate miRNAs that are potentially derived from multiple genomic locations but have identical miRNA sequences, thus cannot be distinguished.

To characterize the tasiRNA composition in the three fractions, we mapped reads to seven *TAS* genes, including three *TAS1*, two *TAS3,* one *TAS2,* and one *TAS4* locus. Just like miRNAs, reads mapping to *TAS* genes accumulated at a significantly lower level in AWF and LSW fractions compared to CL (*SI Appendix,* Fig. S9). Additionally, we also observed that the size distribution of these reads is mainly 21 and 22 nt long in CL, which shifts to 19 and 20 nt in AWF, and 20 and 21 nt in LSW (*SI Appendix,* Fig. S7*B*).

### tRNA Halves Are Enriched in AWF

As noted in Fig. 3*B*, our sRNA sequence analysis revealed that AWF RNA is especially enriched in reads that mapped to tRNA genes. Since tRNA-derived fragments (tRFs) have recently been shown to possess biological activities independent of amino acid delivery to ribosomes (33–35), we analyzed these reads to assess their size distributions, tRNA source, and the start and stop sites of each read. For this analysis, we used the Unitas pipeline (36), which classifies tRNA-derived fragments into 5′ tRFs (5′ end to D-loop), 5′ tR-halves (5′ end to anticodon-loop), 3′ tRFs (TѱC-loop to 3′ end, but without CCA addition), 3′ CCA-tRFs (TѱC-loop to 3’CCA), 3′ tR-halves (anticodon-loop to 3′ end) and misc.-tRFs (miscellaneous tRFs; any reads that map to the mature tRNA but do not align to the very 5’ or the very 3’ ends). The majority of AWF reads were classified in the misc-tRF category (Fig. 5*A*), consistent with a higher level of degradation in AWF samples resulting in the removal of 5’ ends and/or 3’ ends of tRNAs. To discern the genomic origins of these reads, we plotted their length distributions (Fig. 5*B*). This analysis revealed that AWF misc-tRFs had peaks at 17 and 18, 28-30, and 32-35 nt. The first peak corresponds to 5’ and 3’ tRFs that are missing the first or last nucleotide, while the second two peaks likely correspond to tRNA halves (both 5’ and 3’). LSW and CL RNA displayed similar size distributions (*SI Appendix,* Fig. S10 *A* and *B*), but with the first peak shifting to 18 and 19 nt for LSW or 16 and 17 nt for CL. These size distributions are consistent with preferential cleavage of tRNAs in the loop regions rather than stem regions. To account for the possibility that the second two peaks corresponded to tRNA halves lacking the 5’ and 3’ ends of the tRNAs, we re-assigned misc.-tRFs as misc.-5’ tR-halves (if they started within the first four nucleotides of the tRNA and were longer than 28 nt), and to misc.-3’ tR-halves (if they started after position 29 and were longer than 27 nt) (*SI Appendix,* Fig. S10*C*). The majority of AWF misc.-tRFs were reassigned as misc.- 3’ tR-halves, with only a few percent assigned to misc.-5’ tR-halves. Taken together, these results confirm that AWF is enriched in tRNA halves, especially 3’ tR-halves.

**Fig. 5.**
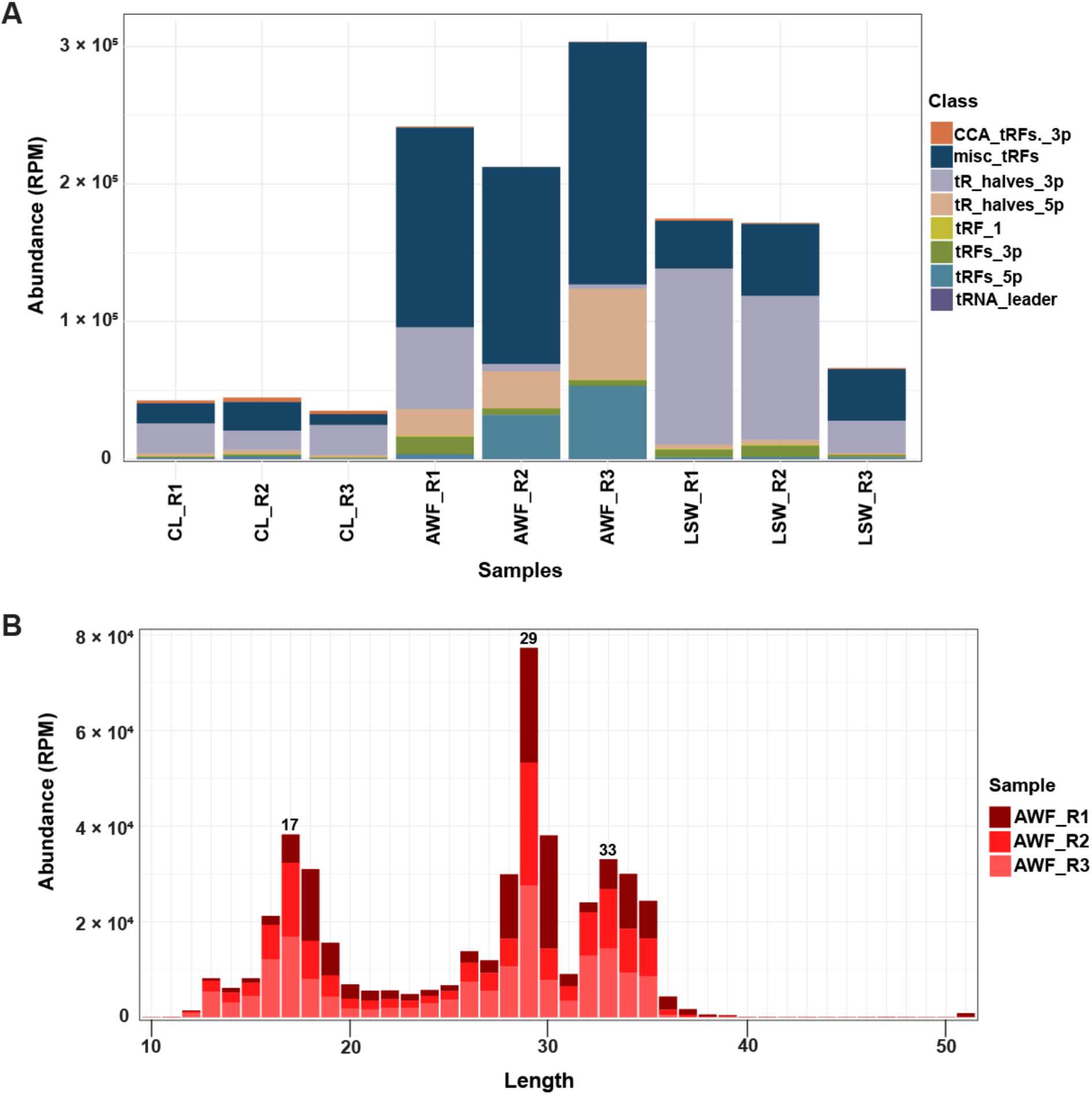
LSW and AWF are enriched in tRNA halves. (*A*) Abundance in reads per million of tRNA-derived small RNAs as classified by unitas (5′-tR-halves, 3′-tR-halves, 5′-tRFs, 3′-tRFs, 3′-CCA-tRFs, tRNA leader and tRF-1) across different samples. (*B*) Length distribution of reads classified as misc-tRF in AWF samples. The colors represent replicates.

We next attempted to confirm the location of the cleavage sites by plotting the start and stop locations of each read that mapped to four of the abundant tRNA families (*SI Appendix,* Fig. S11). This analysis revealed that the vast majority of reads started within the first 10 nucleotides or the anticodon loop (between positions 35 and 45). Notably, the width of the first peak varied significantly between tRNA families, with tRNA^Glu^ being much broader than tRNA^Ala^. Since this was observed across CL, AWF, and LSW samples, it suggests that this is a property of the tRNA, perhaps reflecting the relative resistance of specific tRNAs to exonucleases. Plots of the 3’ position of tRNA reads revealed more variation between tRNA families. For example, tRNA^Ala^ showed clear peaks within the D-loop (positions 14-25), but the second peak was between positions 50 and 60, which corresponds to the TѱC stem, apparently missing 3’ ends within the anticodon loop, despite a clear peak for starts of 3’ tRNAs within this loop. We speculate that this is an artifact of the tRNA sequencing process, with the reverse transcriptase possibly blocked within the D-loop of tRNA^Ala^ sequences, and thus not reaching the anti-codon loop. In contrast to tRNA^Ala^, the sequences derived from tRNA^Glu^, and tRNA^Gly^ family showed a broader peak at 3’ ends that spanned the anti- codon loop and a narrower peak corresponding to 3’ ends within the D-loop. This pattern suggests that the reverse transcriptase has less difficulty traversing the D-loop of these tRNAs.

To evaluate the accumulation of specific tRNA families, we combined all reads originating from tRNAs based on their anticodons. We identified a total of 94 different anticodon tRNAs in our samples, consisting of 50 nuclear, 29 chloroplastic, and 15 mitochondrial tRNAs. Of these 94 tRNAs, a small subset dominated the read count, with tRNA^Glu^_CUC_ and tRNA^Glu^_UUC_ being especially abundant, topping the list for AWF, LSW, and CL (Dataset S1). This abundance likely reflects both expression and resistance to extracellular RNases.

### tRNAs Are Less Processed in LSW than in AWF

The above sequence analyses indicated that AWF tRNAs are more degraded/processed than total CL tRNAs. Processing of tRNA by endoribonucleases in the extracellular environment has previously been reported in mammals. Although tRNA processing in plants has been well studied, whether this occurs primarily in intracellular or extracellular locations has not been assessed. Cleavage of tRNAs by endoribonucleases primarily occurs within tRNA loops (37, 38). The primary enzymes implicated in this processing are endoribonucleases belonging to the RNase T2 family (33). The cleavage of tRNA by T2 RNases has been shown to result mostly in 3ʹ tRNA-derived fragments with 5ʹ-OH ends and 5ʹ fragments with a terminal 2ʹ,3ʹ- cyclic phosphate (cP) or 3’ phosphate (P), both of which are recalcitrant to the sRNA sequence library preparation used in this work. Only a low percentage of tRNA-derived fragments possess the conventional 3ʹ-OH and 5’-P ends, and therefore are likely sequenced (27, 39–41). To overcome these limitations, we selected four tRNAs, tRNA^Gly^, tRNA^Glu^, tRNA^Ala^, and tRNA^Lys^, and analyzed them using RNA gel blot analysis (Fig. 6 and *SI Appendix,* Fig. S12). Three of these tRNAs, tRNA^Gly^_UCC_, tRNA^Glu^_CUC/UUC_, and tRNA^Ala^_AGC_ were selected based on their high abundance in our sRNA sequence data (Dataset S1). tRNA^Lys^_CUU_ was selected because it did not occur in the top 10 list of tRNAs in our sRNA sequence but is known to accumulate stably in mammals without undergoing extensive cleavage (42). To enable the detection of both 5ʹ and 3ʹ tRNA-derived fragments, along with the corresponding full-length tRNAs, we designed two probes for each tRNA, one complementary to the 5’ tR-half (5’ probe) and the other to the 3’ tR-half (3’ probe) (*SI Appendix,* Table S1).

**Fig. 6.**
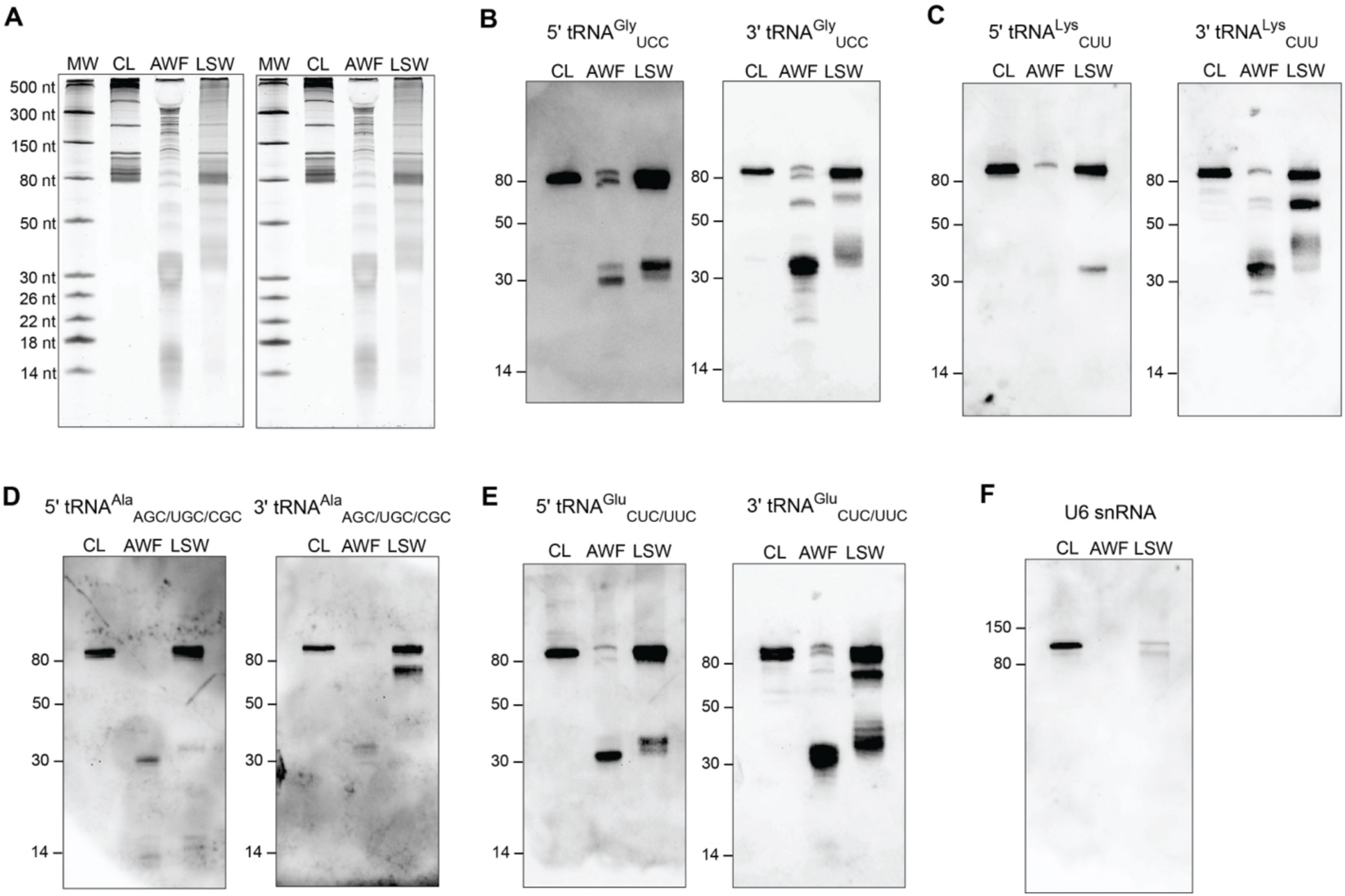
Extracellular fractions are enriched in tRFs compared to whole cell lysate. (*A*) 100 ng of RNA from CL, AWF, and LSW was separated on 15% denaturing polyacrylamide gels and stained with SYBR GOLD nucleic acid stain. (*B*, *C*, *D*, and *E*) Upon blotting onto a positively charged nylon membrane, RNA was probed with DIG-labeled 5’ and 3’ probes against tRNA^Gly^, tRNA^Lys^, tRNA^Ala^, and tRNA^Glu^, respectively, to detect the tRNA halves (30-40 nt), and tiny tRFs (<18 nt) in the CL, AWF, and LSW samples. Full-length tRNAs (>80 nt) and three-quarter (3/4) tRFs (60-70 nt) were also detected in these samples. (*F*) RNA was probed with a DIG-labeled probe against U6 snRNA to assess its presence in AWF and LSW.

These RNA blot analyses confirmed that tRNA halves accumulate to higher levels in AWF than in LSW (*SI Appendix,* Fig. S12 and Fig. 6). Notably, we could not detect any tRNA halves in CL RNA, indicating that cleavage of tRNAs rarely occurs inside plant cells. Interestingly, we found that 3’ tR-halves accumulate even more compared to their 5’ counterparts in both AWF and LSW, suggesting that 3’ tR-halves are more stable than 5’ tR-halves in the extracellular milieu. This was especially noticeable for tRNA^Lys^_CUU_ for which 5’ tR-halves seem to be highly unstable and prone to rapid degradation, while the 3’ tR-halves exhibited stable accumulation in AWF (*SI Appendix,* Fig. S12*C* and Fig. 6*C*). Strikingly, by using the 3’ probes, we were able to discover a class of tRNA-derived fragments ranging in size from ∼60–70 nt in both AWF and LSW, which we termed “tRNA three-quarters” (hereafter referred as tR-3/4). These are likely produced after cleavage only in the D-loop of the full-length tRNA (Fig. 6 *B*, *C*, *D*, and *E*). Moreover, we noted a high accumulation of full-length tRNAs in LSW compared to AWF, suggesting that the tRNAs are much less processed in LSW than in AWF. The occurrence of less tRNA processing in LSW compared to AWF could be attributed to the absence or very low abundance of RNA-processing enzymes on the leaf surface compared to the apoplast (*SI Appendix,* Fig. S6 and Fig. 2 *B* and *C*).

We also observed a differential accumulation of fragments derived from the same parental tRNAs but with slightly different lengths. In mammals, tRNA^Gly^ has been shown to undergo sequential cleavage, first cleaved at the anticodon loop, generating 34 and 35 nt 5′ tR-halves that rapidly disappear, which are subsequently replaced by highly stable shorter fragments of approximately 30 and 31 nt (42). We observed a higher accumulation of 34 and 35 nt 5’ tR-halves of tRNA^Gly^ in LSW, and stable accumulation of the corresponding 30 and 31 nt 5’ tR-halves in AWF. We also observed a similar trend upon specific assessment of tRNA halves derived from tRNA^Gly^ in our sequencing data. This further substantiates our hypothesis that there is less processing of RNA in LSW compared to AWF.

To assess whether the secretion of full-length tRNAs in the extracellular spaces is a selective process, we used a probe specific to the U6 small nuclear RNA (snRNA), an essential component of the catalytic core of the pre-mRNA processing spliceosome (Fig. 6*F*). U6-specific probes have previously been used as a loading control in northern blot analyses of small RNAs present in whole CL samples. We detected little to no U6 RNA in exRNA derived from LSW and AWF fractions, suggesting that U6 is either not secreted or secreted and is highly sensitive to extracellular RNases in AWF.

### AWF and LSW exRNAs Display Reduced Abundance of Many Transcripts Relative to Total CL RNA

To investigate the long RNA content of AWF and LSW, we used the above described samples and generated standard RNA sequencing libraries without employing an rRNA depletion step. We used the standard Tuxedo pipeline (43) and the Arabidopsis TAIR10 annotation to identify differences in transcript abundance between the three fractions, CL, AWF, and LSW. In comparing AWF and LSW to CL, we found 300 and 297 genes, respectively, that were differentially expressed (DE; Fig. 7*A*, and Dataset S2). Additionally, we found 176 DE genes when comparing AWF to LSW (Fig. 7*A*). To understand the gene overlap between AWF and LSW, we examined the DE genes common to both extracellular fractions. Most DE genes were unique either in LSW when compared to CL (172 genes) or in AWF when compared to CL (158 genes) (Fig. 7*C*). The majority of these unique DE genes were downregulated, demonstrating that both AWF and LSW fractions are gene-depleted relative to total cellular RNA. Interestingly, the seven genes with higher accumulation in LSW correspond to four pre-tRNAs, two transposable elements, and a GTP binding Elongation factor Tu family protein. The remaining 165 genes, which have lower accumulation levels in LSW, are enriched in GO terms related to response to stimulus (Dataset S3). Conversely, the 25 genes with higher accumulation in AWF compared to CL have a GO enrichment in the biotic stress category. The other genes with lower accumulation in AWF are enriched in GO terms related to the photosynthetic electron transport chain, seed development, and stress responses. Additionally, we observed that 61 genes were uniquely differentially expressed in AWF compared to LSW (Fig. 7*C*), all of them with lower abundance in AWF. These genes are enriched in GO categories related to both biotic and abiotic stress responses. This supports the hypothesis that AWF and LSW exhibit different RNA accumulation patterns and that the role of RNAs in these fractions is linked to both biotic and abiotic responses.

**Fig. 7.**
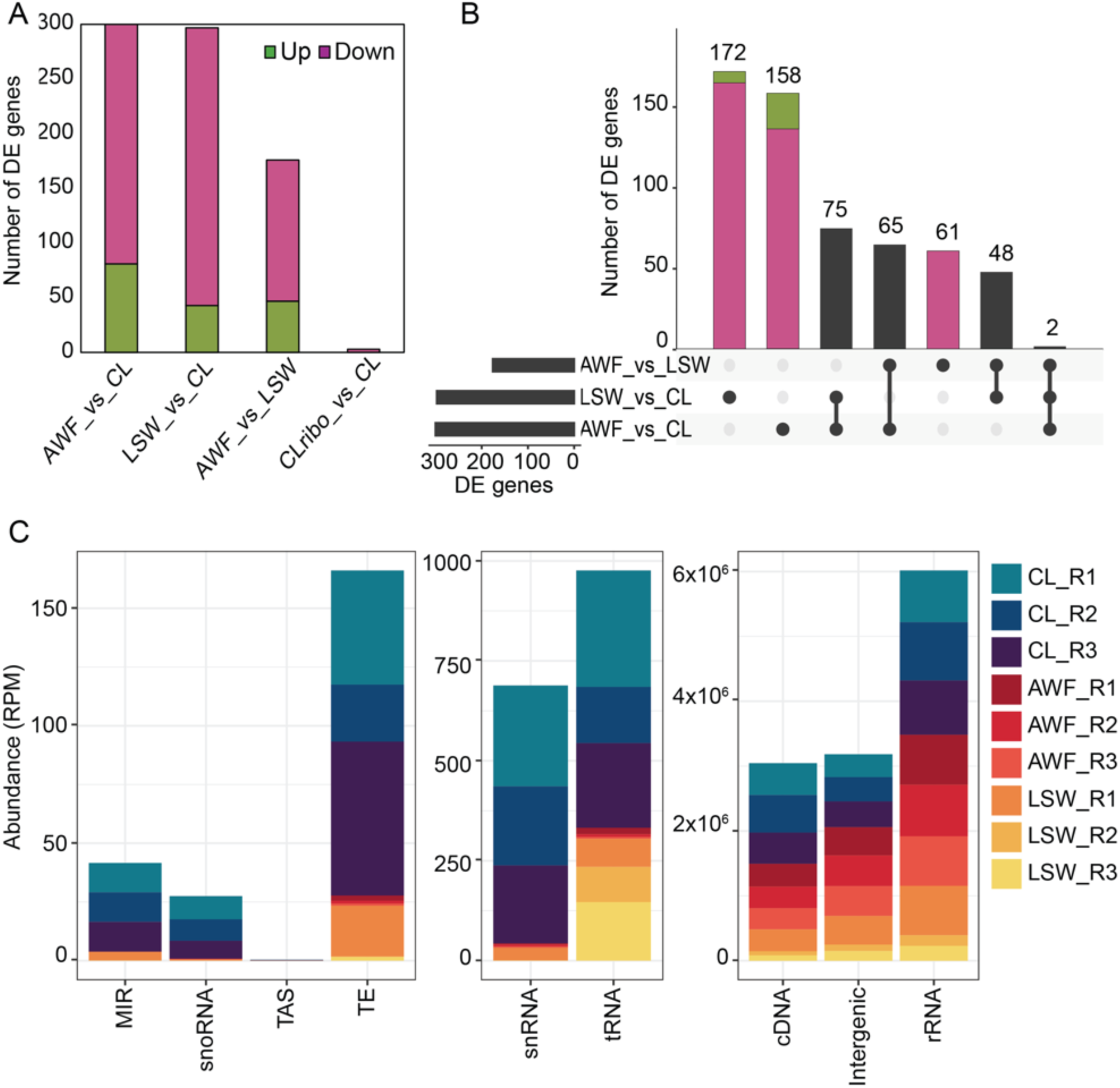
AWF and LSW RNAs are depleted in protein-coding transcripts. (*A*) Number of differentially expressed genes found in the comparison of each fraction, down-regulated in pink and up-regulated in green. (*B*) Shared and unique DE genes in each comparison, the bottom left plot shows the entire size of each set as a horizontal histogram, the bottom right shows the intersection matrix and the upper right shows the size of each combination as a vertical histogram, with down-regulated genes colored in pink and up-regulated genes colored in green. (*C*) Genomic origin of RNA reads based on the categories established in the TAIR 10 genome version. RNAs that mapped to the genome were categorized by origin and plotted by relative abundance in RPM. The x-axis represents the feature of interest. Note the differing Y axis values between the three panels. MIR, microRNA precursors; snoRNA, small nucleolar RNA; TAS, trans-acting siRNA precursors; TE, Transposable elements; snRNA, small nuclear RNAs; tRNA, transfer RNAs; cDNA, complementary DNA; Intergenic regions; rRNA, ribosomal RNAs; The y-axis indicates the relative cumulative abundance in RPM, calculated as indicated before. The three colored shades represent three independent biological replicates, with each replicate derived from 18 Arabidopsis plants.

The remaining DE genes were categorized into four groups based on the number of comparisons where they showed a statistically significant difference (Dataset S2). The first group contains 75 genes commonly DE in both AWF and LSW compared to CL. Of these, only five (5) genes show higher accumulation in LSW and AWF compared to CL, including four rRNA genes and a transposable element. The other 70 genes show lower accumulation in AWF and LSW compared to CL and are mainly associated with water deprivation according to their GO enrichment. The second group includes 65 genes that are DE in AWF compared to both CL and LSW. Sixteen of these genes show lower accumulation in AWF compared to CL, but higher accumulation in AWF compared to LSW. Conversely, 49 genes have higher accumulation in AWF compared to CL and lower accumulation in AWF compared to LSW. This group is enriched in GO terms related to plant defenses, including oxylipin metabolism. The third group contains 48 genes that are statistically significant in LSW compared to both CL and AWF. Seventeen of these have a lower abundance in LSW compared to both CL and AWF, while 31 genes have a higher accumulation in LSW. The final group includes only two genes common to all comparisons: a GATA transcription factor (AT3G54810) and other RNA (AT1G70185).

Broadly speaking, genes that accumulate significantly more in AWF compared to CL are enriched in GO terms related to biotic stress. Conversely, genes with significantly less accumulation in AWF compared to CL are enriched in GO terms for stomata movement, photosynthesis, water deprivation, and biotic stresses. In the case of LSW, genes with significantly more accumulation compared to CL are enriched in terms of iron transport, translation, ATP biosynthetic processes, and aerobic respiration. Genes with significantly less accumulation in LSW compared to CL are enriched in GO terms for water deprivation, response to temperature stimulus, and signal transduction. These accumulation patterns suggest that the RNA found in AWF and LSW might have different origins, or there is differential degradation occurring in AWF compared to LSW, which would be consistent with the tRNA analyses described above.

We also analyzed the origin of the RNAs captured in each fraction (Fig. 7C). We observed that all fractions have similar levels of RNAs from rRNA, cDNA, and intergenic regions and that these are the most abundant ones. Interestingly, long reads originating from tRNAs were highly enriched in LSW compared to AWF, which is consistent with the RNA gel blot analyses that showed a marked reduction in full-length tRNAs in AWF. Notably, the exRNA fractions were depleted in reads originating from all the other minor features, including miRNA precursors, suggesting that these are particularly unstable in the extracellular environment.

### exRNAs May Form Cation-Dependent Aggregates or Condensates *In Vivo*

In our previous work, we found that apoplastic RNAs longer than 50 nt could be pelleted by ultracentrifugation at 100,000 *g*, indicating that this RNA was associated with some kind of particle. We thus assessed whether RNAs found on the leaf surface were also associated with a particle of some kind that could contribute to their stability. We ultracentrifuged AWF and LSW at 40,000 *g* (P40), and then the supernatant of this pellet at 100,000 *g* (P100-P40). These speeds are commonly used to isolate different subpopulations of plant EVs, but we did not expect single RNA molecules to pellet even at 100,000 *g*. We then analyzed the RNA content of P40 and P100-P40 pellets along with the supernatant of the P100-P40 pellet (S100). In concordance with our prior work (11), apoplastic RNAs longer than 35-40 nt pelleted at both speeds, although a higher amount was observed at 100,000 *g* (Fig. 8*A*). Surprisingly, despite the lack of proteins in LSW, long LSW RNAs also pelleted at both speeds (Fig. 8*A*).

**Figure 8.**
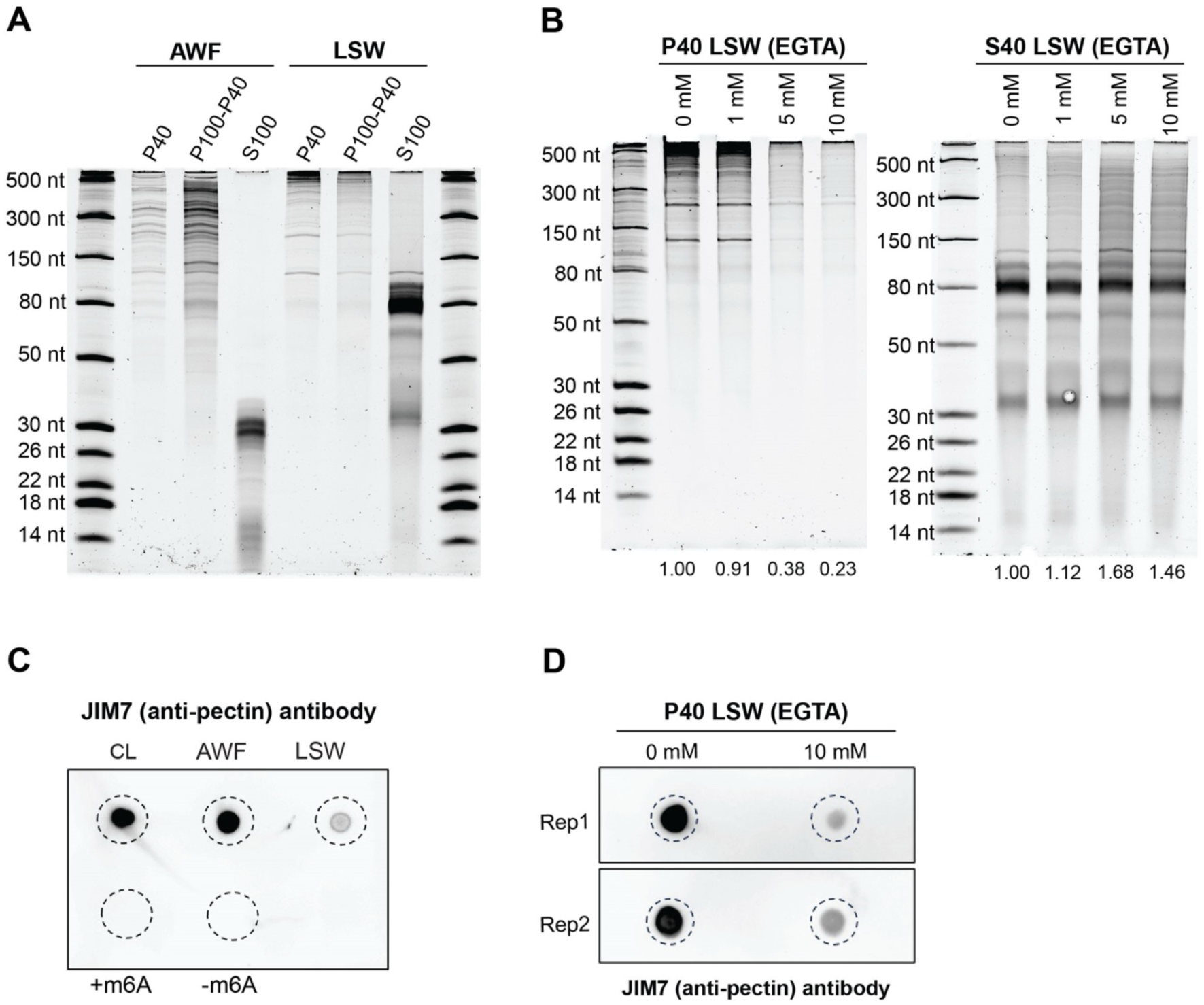
Long RNAs in LSW form cation-dependent aggregates or condensates. (*A*) Profile of RNAs in P40, P100-P40 pellets, and supernatants (S100) obtained using AWF and LSW as the starting material. Volumes of starting AWF and LSW were normalized by fresh weight. RNAs were extracted using TRIzol, separated on a 15% denaturing polyacrylamide gel and stained with SYBR Gold nucleic acid stain. (*B*) Cations contribute to RNA particle formation. LSW was treated with increasing concentrations of EGTA as indicated for 20 min on ice, followed by ultracentrifugation at 40,000 *g*. RNAs were isolated from P40 pellets and their corresponding supernatants (S40) using TRIzol, separated on a 15% denaturing polyacrylamide gel and stained with SYBR Gold nucleic acid stain. RNA abundance in each gel lane was estimated by densitometry and expressed relative to the negative control. Experiments A and B were repeated three times with similar results. *(C)* TRIzol-purified RNA contains pectin. An aliquot of 100 ng of total CL, AWF, and LSW RNA was dot-blotted onto a positively charged nylon membrane and then probed with an anti-pectin antibody (JIM7). For negative controls, 100 ng of synthetic 21 nt RNAs with identical sequences (except for a single m6A modification on +m6A oligo) were used. *(D)* Cations contribute to co-pelleting of pectin with LSW RNA. LSW was treated with 0 and 10 mM EGTA for 20 min on ice, followed by ultracentrifugation at 40,000 *g*. RNAs were isolated from P40 pellets and aliquots of 100 ng of RNA were dot-blotted onto a positively charged nylon membrane and then probed with the JIM7 (anti-pectin) antibody.

Considering that RNAs are negatively-charged molecules, we then tested whether cations were involved in RNA pelleting. Divalent ions can bind directly to more than one phosphate group and form a bridge between different parts of an RNA molecule or between different RNA molecules (44). Since we used a buffer containing 2 mM CaCl_2_ to isolate LSW, we repeated the pelleting at 40,000 *g* (P40) after adding increasing concentrations of EGTA, a divalent cation chelator with a strong affinity for Ca^2+^, to the LSW. We observed a significant reduction in the pelleting of RNA with 5 and 10 mM of EGTA (Fig. 8*B* and *SI Appendix,* Fig. S13*A*); additionally, we eliminated Ca^2+^ from VIB and repeated the LSW isolation and pelleting, but no effect was observed in the amount of RNA that pelleted compared to the buffer containing Ca^2+^ (*SI Appendix,* Fig. S13*B*), suggesting that leaf surfaces possess a substantial amount of endogenously secreted Ca^2+^ or other cations that facilitate RNA aggregation or condensation on the leaf surface.

The promotion of RNA pelleting from LSW by cations suggested that there could be other negatively charged molecules associated with the RNA. An abundant negatively charged molecule secreted by plant cells is pectin, a cell wall polysaccharide that can be non-covalently crosslinked by Ca^2+^ ions. To assess whether pectin was present in P40 pellets, we use an anti-pectin antibody (JIM7). Since pectin is highly methylated, we also used 21 nt long oligonucleotides with and without a single modified methylated adenosine as negative controls to assess the specificity of JIM7. We confirmed the presence of pectin in RNA isolated from the CL, AWF, and LSW fractions (Fig. 8*C*). We next assessed whether pectin precipitation was also inhibited by EGTA, similar to RNA. We observed a significant reduction in pectin in the P40 pellet in the presence of 10 mM EGTA compared to no EGTA (Fig. 8*D*), suggesting that pectin associates with RNA and could potentially stabilize RNA on the leaf surface.

### exRNA Is Not Enriched in m^6^A Modification

Previously, we reported that exRNA isolated from AWF is enriched in the post-transcriptional RNA modification N6-methyl adenine (m6A). To test whether exRNA isolated from LSW is also enriched in m6A, we performed a dot blot analysis using an m6A-specific antibody as described previously (11). This analysis indicated that LSW RNA was highly enriched in m6A modification relative to both total cellular RNA and AWF (Fig. 9*A*). We questioned this result, however, as the signal was much stronger than our positive control, which is a 21 nt oligonucleotide containing a single modified adenine. We thus hypothesized that the m6A antibody used in this assay might be binding to a contaminating molecule that co-purified with RNA. We used two different sources of m6A antibodies to perform the dot blot analysis and obtained similar results, indicating that this potential cross-reactivity was not unique to a single antibody source. In addition, the m6A signal in AWF RNA samples treated with two different RNases remained comparable to untreated samples, despite a significant reduction in total RNA amount. (*SI Appendix,* Fig. S14 *A* and *B*). This observation strengthened our hypothesis that commercial anti-m6A antibodies cross-react with a non-RNA molecule that co-purifies with RNA extracted from leaf-derived samples.

**Fig. 9.**
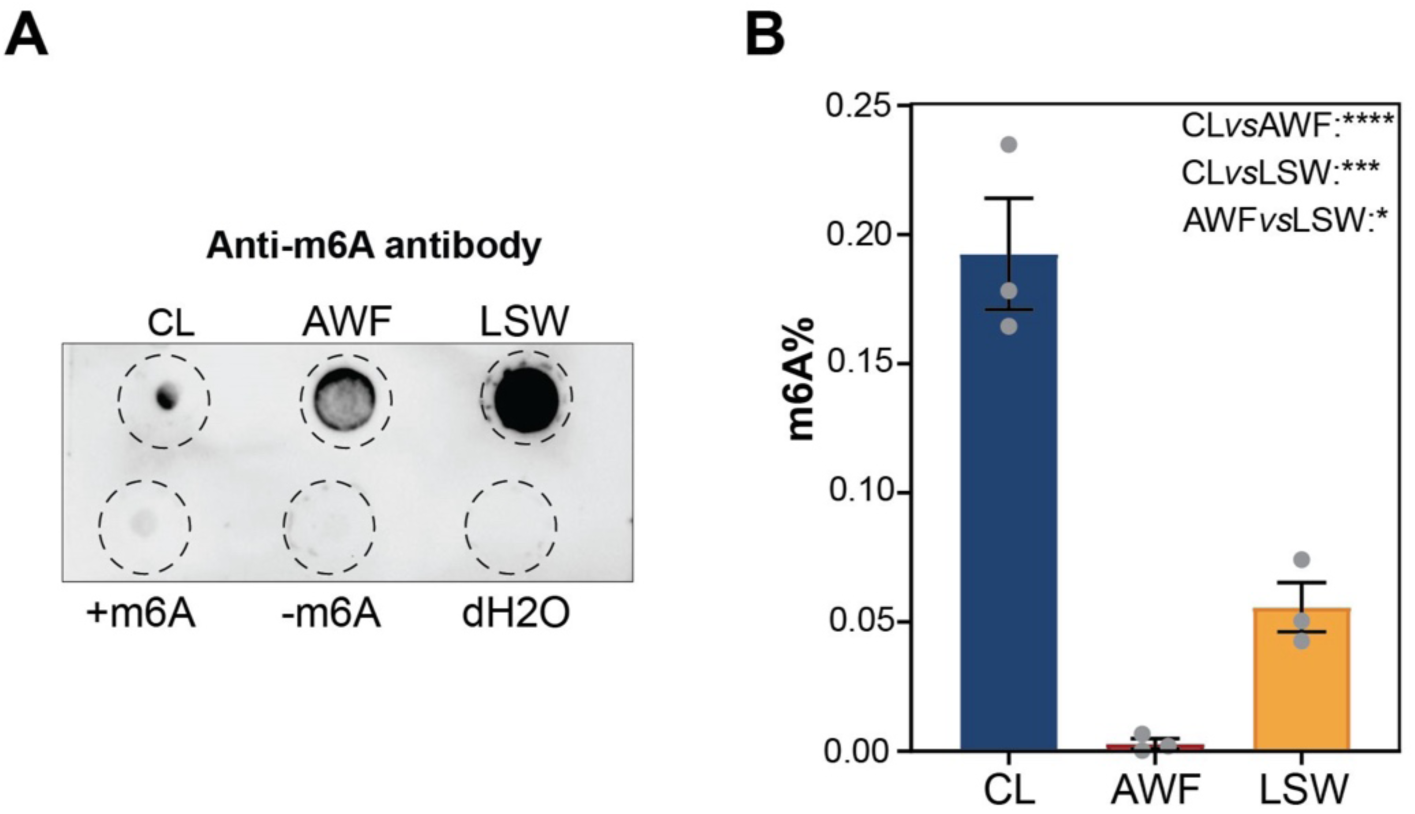
**exRNA is not enriched in m6A modification. (***A*) Dot blot analysis of m6A content. An aliquot of 100 ng of total CL, AWF, and LSW RNA was dot-blotted onto a positively charged nylon and then probed with the anti-m6A antibody. For positive and negative controls, 100 ng of synthetic 21-nt RNAs with identical sequences except for a single m6A modification on the positive control were used. (*B*) ELISA-based analysis of m6A content. Bar graph indicates the percentage of m6A modification in each sample assessed using an RNA-specific ELISA-based m6A modification kit. The values are calculated using a positive control with 100% m6A modification. Significant differences were calculated using one-way ANOVA and Fisher’ s LSD post-hoc test ****, P<0.0001; ***, P<0.001; *, P<0.05.

Due to this concern, we pursued alternative methods of quantifying m6A content. Specifically, we tested an ELISA-based method that has recently been commercialized (45). In this method, RNA is first bound to the wells of microtiter plates and extensively washed prior to incubation with the m6A detection antibody, which we hoped would remove any contaminating molecules. This method allowed us to determine the percentage of m6A modification in total CL, AWF, and LSW RNA. These analyses revealed that exRNA isolated from AWF is not enriched but depleted in m6A modification compared to total CL RNA (Fig. 9*B*), which is contrary to what we observed with the dot blot analysis (Fig. 9*A*). However, LSW RNA seems to be enriched in m6A modification compared to AWF RNA, while being about three-fold lower than total CL RNA (Fig. 9*B*). This result indicates that plant exRNA is not enriched in m6A modification.

## Discussion

In our previous work, we reported that the plant leaf apoplast contains abundant and diverse species of RNA outside EVs, which are protected from RNase degradation by a group of extra-vesicular RNA-binding proteins (10, 11). These apoplastic RNAs have not been ascribed a biological function, but potential roles have been discussed in a recent review (1). Here we have reported for the first time the existence of abundant plant RNAs on the leaf surface. The RNA content on the leaf surface is distinct from that of the apoplast, as it appears much less degraded, which suggests an absence of RNA processing enzymes on the leaf surface. Additionally, our RNase protection analyses indicated that although leaf surface RNAs are less processed than apoplastic RNAs, they are not protected by proteins or vesicles. Consistent with an absence of RNases on the leaf surface, we found very low amounts of protein overall in LSW samples.

In our previous reports, we and others demonstrated that most siRNAs that co-pellet with EVs are located outside of EVs and can be found associated with proteins (10, 11, 22). Moreover, the vast majority of sRNAs in the apoplast did not pellet, even after spinning at 100,000 *g* (11), a speed that has been extensively used to isolate sRNA-containing EVs (18, 20). Those results encouraged us to analyze the entire sRNA pool in the extracellular space without conducting any ultracentrifugation steps. Surprisingly, our results revealed that Arabidopsis does not seem to accumulate siRNAs/miRNAs in extracellular fractions, with most miRNAs identified by sRNA sequencing far more abundant inside cells than outside cells when quantified in terms of reads per million.

The most abundant class of exRNAs that we found in the apoplast was tRNA-derived fragments (tRFs) and tRNA halves (collectively referred to as tRNA-derived RNAs (tDRs). In mammals, full-length tRNAs are secreted into the extracellular space from diverse cell types, where they are rapidly processed into tDRs by extracellular RNases (8, 9). Although this mechanism has not been explored in plants, we believe that the extracellular processing of tRNA also occurs in plants. The generation of tDRs in plants relies mainly on endonucleases belonging to the RNase T2 family (reviewed in 33). Of particular relevance to our findings, these RNases contain signal peptides and are known to be secreted to the apoplast. Consistent with this, the ratio of tRNA halves to full-length tRNAs in the apoplast was very high, while the opposite was observed in RNA isolated from total cell lysate, suggesting a lack of processing of tRNAs inside the cell. Although the pattern observed could also indicate selective and rapid secretion of tDRs by cells, we believe it is far more likely that tDRs are generated outside the cell due to the presence of extracellular RNases in the apoplast.

Given the presence of RNases in the apoplast, it is somewhat surprising that single-stranded sRNAs such as tDRs are so stable in the apoplast. In mammals, it has been shown that some tDRs can acquire high resistance to RNases by forming self-protecting homo- and hetero- dimers or oligomers (46, 47). Moreover, a recent mammalian paper shows that after cleavage at the anticodon loop, the 5’ and 3’ halves of the original tRNA can be held together, thus stabilizing these tDRs (42). Whether similar processes contribute to tDR stability in plants is still unknown. Interestingly, according to our RNA gel blot analyses, 3’ tDRs are more stable than 5’ tDRs in the apoplast. We speculate that posttranscriptional modifications may help stabilize specific tDRs (48, 49). Alternatively, 3’ tDRs may fold into more stable secondary structures than 5’ tDRs.

We have also shown the existence of 3/4 length 3’ tRFs. These tRFs are highly abundant in the apoplast and are likely produced after cleavage in the D-loop. Of note, we did not observe any ¾ length 5’ tRFs (produced by cleavage in the TѱC loop), and we detected very few 3’ tRFs. This suggests that cleavage in the TѱC is less common than in the two other loops. In addition, our results support the idea that cleavage in the TѱC takes place only after cleavage in the anticodon loop.

Diverse biological functions of tDRs have been recently proposed in plants, including a role in regulating gene expression (reviewed in 33, 34). Although direct evidence of a functional role for extracellular tDRs in cross-kingdom gene regulation is lacking, the high accumulation of tDRs in the apoplast suggests that they are biologically relevant. A key question to be addressed in the future is whether plant cells can take up tDRs from the apoplast, which would indicate that tDRs could function in intercellular gene regulation. Another key question is whether plant tDRs are taken up by leaf bacteria and can impact bacterial growth. Recent work on the human oral microbiome suggests that this is likely (50, 51). In that work, it was shown that human saliva contains abundant tDRs, several of which bear similarity to tRNA sequences in specific oral bacterial species. Application of these tDRs to cultures of these bacterial species inhibited the growth of some species, and not others, and this inhibition was sequence specific. Thus, human tDRs help structure the human oral microbiome. Notably, this effect was observed with naked RNA, thus association with protein or packaging inside EVs was not required for uptake of tDRs by bacteria.

Our discovery that leaf surfaces are coated with RNA raises the question of how this RNA gets to the surface and whether it is being secreted directly from epidermal cells, passing through the cuticle, or instead, is secreted from mesophyll cells and passes through stomata. If the latter were the case, however, we would expect LSW RNA to look more like apoplastic RNA. Instead, we observed that LSW RNA appears much less degraded than apoplastic RNA, indicating that a mesophyll origin is unlikely. How RNA molecules pass through the hydrophobic cuticle layer and reach the leaf surface is unknown, but the lack of RNA-binding proteins in LSW makes it unlikely that RBPs function as RNA-carriers across the cuticle. We speculate that regions of higher permeability across the cuticle layer can facilitate RNA diffusion. For instance, the cuticle covering trichomes is known to be more permeable than the cuticle of pavement cells in tomato (52).

It is also possible that our isolation protocol partially extracts RNA from the cuticle. Leaf surface RNAs may be embedded inside the cuticular layer along with other high-molecular-weight polysaccharides, such as pectins, which are soluble in aqueous and acidic conditions, and can be non-covalently crosslinked by Ca^2+^ ions (53). During LSW extraction, we use a slightly acidic VIB buffer that contains Ca^2+^ along with a wetting agent (Silwet). These properties could facilitate the release of RNAs and pectin from the cuticular layer and while promoting their association with each other.

In support of the latter suggestion, RNA pelleting was highly reduced when LSW was pretreated with EGTA before ultracentrifugation, suggesting that aggregation of RNAs in LSW is mediated by Ca^2+^ and other divalent cations. We speculate that the presence of polygalacturonic acid/pectins on the leaf surface might contribute to RNA aggregation. These polysaccharide molecules are heavily negatively charged like RNA and might be involved in the formation of a complex mesh with RNA and Ca^2+^ leading to the formation of a gel-like substance.

In the quest to identify factors associated with the selective secretion and stabilization of leaf surface RNA, we also tested if the secreted LSW RNA exhibits a higher level of posttranscriptional modification compared to AWF and CL RNA. Using an ELISA-based assay, we found that neither AWF nor LSW RNA is enriched in m6A modification relative to the total CL RNA. Therefore, it will be necessary to test for the enrichment of other modifications, which might provide insights into the secretion and stability of exRNA.

Our discovery of extravesicular RNA on leaf surfaces reinforces our previous finding that extravesicular RNAs are main constituents of the exRNA pool in plants. Our work highlights the need to identify alternate mechanisms through which RNA might be secreted and trafficked independent of EVs. This surprising finding of RNA on the leaf surface opens important questions about the mechanisms involved and its function. For instance: What are the cellular sources of RNA on the leaf surface and in the apoplast? How does this RNA pass through the plasma membrane and cell wall? Does leaf surface RNA play a role in shaping the leaf microbiome?

Exploring these questions along with the potential roles of the secreted RNA in plant-microbe interaction will provide valuable information for improving crop plant protection strategies, especially in the context of improving the efficacy of HIGS and SIGS.

## Materials and Methods

### Plant Materials and Growth Conditions

*Arabidopsis thaliana* Col-0 wild type (WT) seeds were grown in 4-inch round plastic pots containing Sungro Propagation Mix. After sowing, the seeds were watered with RootShield Plus WP Biological Fungicide (600 mg/L, catalog no. 68539-9, BioWorks), covered with a clear plastic dome, and kept at 4 °C in darkness to induce synchronous germination. After 3 days, the pots with seeds were transferred to a temperature-controlled short-day growth chamber set at 24 °C under a 9 h light/15 h dark photoperiod with 150 µmol m^−2^ s^−1^ photosynthetic photon flux density (50:50 mix of 3,500 and 5,000 K spectrum GE HI-LUMEN XL Starcoat 32-watt fluorescent bulbs). After 10 days, individual seedlings were transferred to 36-cell tray inserts containing Sungro Professional Growing Mix and covered with a clear plastic dome for one to two weeks. Miracle-Gro water-soluble fertilizer (catalog no. 24-8-16) was used (250 mg/L) to water the plants every 10-15 days. The GRP7-GFP transgenic line was a gift from Prof. Dr. Dorothee Staiger at Bielefeld University, Bielefeld, Germany. This line expresses a genomic copy of GRP7 fused to GFP, including the GRP7 5’-UTR, intron, and 3’-UTR under the control of the native GRP7 promoter in the *grp7–1* mutant background (54).

### Isolation of Leaf Surface Wash (LSW) and Apoplastic Wash Fluid (AWF)

For each replicate, leaf surface wash (LSW) and apoplastic wash fluid (AWF) were isolated from six-to-seven-week-old Arabidopsis plants as depicted in Supplementary Figure 1. For LSW isolation, vesicle isolation buffer (VIB; 20 mM 2-(N-morpholino) ethanesulfonic acid pH 6.0, 2 mM CaCl_2_ and 0.01 M NaCl) supplemented with 0.001% (v/v) of Silwet-77 (Phytotech Labs, Product ID S7777) (a rapid wetting agent that promotes low surface tension, better adhesion, and coverage on foliar surfaces) was sprayed on both sides of the detached whole rosettes. To recover LSW, the sprayed rosettes were carefully placed inside needleless 60 mL syringes containing holes at the bottom (two rosettes per syringe) placed inside 250 mL centrifuge bottles and centrifuged for 10 min at 100 *g* at 4 °C (JA-14 rotor, Avanti J-20 XP Centrifuge; Beckman Coulter, Indianapolis, IN, USA). LSW was then filtered through a 0.2 µm syringe filter (Acrodisc syringe filter, Pall Corporation, New York, USA). Thereafter, the same set of plants was washed with distilled water and used to isolate AWF following the protocol described previously (21) with minor modifications. Briefly, rosettes were vacuum infiltrated for 20 sec with VIB. After vacuum infiltration, excess buffer was removed from the leaf surfaces by gentle blotting with Kimwipes. To collect the AWF, rosettes were placed inside 60 mL needleless syringes as described for LSW collection and centrifuged at 4 °C for 30 min at 600 *g* with slow acceleration. AWF was then filtered through a 0.2 µm syringe filter. Filtered LSW and AWF were either used immediately or stored at –80 °C until further use. The fresh weight (FW) of the plants used for each replicate was noted and subsequently used to estimate the amount of RNA and proteins per gram FW. For all experiments, a biological replicate was considered as the batch of a given number of plants growing in the same 36-cell insert that were sown at least one week apart from the other biological replicates.

### Isolation of Particles from AWF and LSW

To obtain pellets containing extracellular vesicles and other particles, freshly isolated AWF and LSW was transferred to ultracentrifuge (UC) tubes and centrifuged at 40,000 *g* for 1 h at 4 °C (TLA100.3 rotor, Optima TLX Ultracentrifuge; Beckman Coulter). Where indicated, EGTA (Ethylene glycol- bis (β-aminoethyl ether)-N,N,Nʹ,Nʹ-tetraacetic acid; Sigma-Aldrich, USA, Product ID E4378) was added to the LSW at the specified concentration, mixed, and incubated for 20 min on ice before the ultracentrifugation step. The supernatant of the P40 pellet was recovered for analysis. To obtain P100-P40 pellets, the supernatant after the 40,000 *g* spin was transferred to ultracentrifuge tubes and centrifuged at 100,000 *g* for 1 h at 4 °C (TLA100.3 rotor, Optima TLX Ultracentrifuge; Beckman Coulter). P40 and P100-P40 pellets were resuspended in cold and filtered VIB or 100 mM Tris pH 7.4 and either used immediately or stored at –80 °C until further use.

### Quantification of Cell Rupture Using Trypan Blue Staining

Leaves were harvested from three individual six-to-seven-week-old Arabidopsis plants before and after LSW and AWF isolation. For staining, a stock solution of trypan blue (10 mL liquified phenol, 10 mL lactic acid, 10 mL glycerol, 10 mL de-ionized water, and 0.02 g of trypan blue (Sigma-Aldrich- 302643-25G)) was prepared, which was then diluted with 95% ethanol (1:2 v/v) as a working solution. Samples were immersed in trypan blue working solution, boiled for 1 min, and then incubated overnight with gentle shaking. For destaining, a chloral hydrate solution was prepared by mixing 1000 g of chloral hydrate (Sigma- Aldrich- 302-17-0) in 400 mL de-ionized water. The stained leaves were incubated in chloral hydrate solution overnight and the solution was replaced once or twice. The leaves were mounted on glass slides with 25% glycerol solution and imaged using a light microscope.

### RNA Extraction

Total leaf RNA (cell lysate) was isolated from 100 mg of leaf tissue using TRIzol Reagent (Thermo Fisher Scientific™, Waltham, MA, USA). Briefly, leaf tissue was frozen in liquid nitrogen and ground into powder using a mortar and pestle. The powder was placed in 1.5 mL centrifuge tubes and 1 mL of TRIzol was added to each tube and mixed by vortexing. The tubes were placed on a tabletop rotator for 10 min at room temperature (RT). Thereafter, 200 µL of chloroform was added to each tube, followed by a brief but vigorous vortexing step. Tubes were allowed to stand at RT for about 3 min, and then centrifuged at 13,000 *g* for 15 min at 4 °C. The aqueous phase was transferred to labeled 1.5 mL centrifuge tubes containing 10 µg of RNase-free glycogen, mixed with one volume of cold isopropanol, and incubated for no more than 1 h at –20 °C. To pellet the RNA, the tubes were centrifuged at 13,000 *g* for 20 min at 4 °C. RNA pellets were washed twice using ice-cold 70% EtOH and resuspended in 20–30 µL ultrapure DNase/RNase-free water. To isolate RNA from P40 and P100-P40 pellets, 1 mL of TRIzol was added to 100 µL of resuspended pellets, followed by the same procedure as described for the total leaf RNA isolation.

To isolate RNA either from supernatant, AWF, or LSW, the RNA was first precipitated by mixing the required volume of supernatant, AWF, or LSW with 0.1 volume of 3 M sodium acetate (pH 5.2) and 1.0 volume of cold isopropanol, incubated at –20 °C for a minimum of 1 h to overnight, and then centrifuged at 13,000 *g* for 30 min at 4 °C. The pellets were washed twice with ice-cold 70% EtOH, resuspended in 100 µL of ultrapure DNase/RNase-free water (Invitrogen™, Waltham, MA, USA), and transferred to 1.5 ml centrifuge tubes. Thereafter, 1 mL of TRIzol was added to each tube and RNA extraction was performed following the same procedure as previously described for the total leaf RNA. Finally, RNA pellets were resuspended in 10 µL of ultrapure DNase/RNase-free water and stored at –80 °C. If needed, to remove phenol or guanidine contamination from the RNA samples, one or two serial precipitations with EtOH and ammonium acetate were performed. Briefly, 1 µg of glycogen was added to 10 µL of RNA and mixed with 0.5 volume of 7.5 M ammonium acetate prior to the addition of 2.5 volumes of 100% EtOH. The mixture was incubated at –80 °C for 30 min and centrifuged at 13,000 *g* for 20 min at 4 °C. The RNA pellet was washed with 70% EtOH and resuspended in 10 µL of ultrapure DNase/RNase-free water. Nanodrop analysis was performed to determine RNA concentrations, which were confirmed using densitometry analysis of RNAs separated on polyacrylamide gels (see below).

### Ribonuclease Protection Assays

To assess whether RNA in AWF and LSW was protected from ribonuclease digestion by either encapsulation inside of EVs or association with proteins, we followed our previously described protocol (11) with minor modifications. Freshly isolated LSW, AWF, P40, or P100-P40 pellets were split into 4 aliquots. Aliquot #1 was treated with 1% (v/v) Triton X-100 in 100 mM Tris pH 7.5 for 40 min with gentle shaking at RT to disrupt EVs. The other three aliquots (aliquots #2, #3, and #4) were suspended in 100 mM Tris pH 7.5 and incubated on ice. Aliquots #1 and #2 were then treated with 1 mg/mL trypsin (Promega, Madison, WI, USA) and incubated at 37 °C for 1 h followed by the addition of 1.5 mg/mL trypsin inhibitor (Worthington Biochemical Corp, Lakewood, NJ USA) to inactivate trypsin. The remaining two aliquots (aliquots #3 and #4) were also suspended in 100 mM Tris pH 7.5 with aliquot #3 incubated at 37 °C and aliquot #4 was placed on ice. Then, DNase and protease-free RNase A (Thermo Scientific™, catalog no. EN0531; diluted in 15 mM NaCl, 10 mM Tris–HCl pH 7.5) was added to aliquots #1, #2, and #3 at a final concentration of 9 µg/mL and the aliquots incubated for 1 h at RT. To inhibit RNase A activity, a mixture of RNase Inhibitor, Murine (APExBIO) and RNase Out (Invitrogen) was added to the aliquots. The four aliquots were stored at –80 °C until analysis.

Additionally, to assess the level of protection of AWF RNAs by proteins, AWF samples were suspended in 15 mM NaCl, 10 mM Tris-HCl pH 7.5 and treated with DNase and protease-free RNase A (Thermo Fisher Scientific™, catalog no. EN0531) at a final concentration of 0, 5, 9, or 20 µg/mL and incubated for 1 h at RT.

### Denaturing Polyacrylamide Gel Electrophoresis of RNAs

Mini gels (7.2 cm x 8.6 cm x 0.75 mm) containing 10% or 15% polyacrylamide and 7 M urea in 1X Tris-Boric Acid EDTA (TBE, pH 8.4) were made using 40% Acrylamide/Bis Solution, 37.5:1 (Bio-Rad, catalog no. 1610148). RNA samples were mixed (1:1) with 2X denaturing loading buffer (95% formamide, 10 mM EDTA, 0.02% SDS, 0.02% bromophenol blue, and 0.01% xylene cyanol), denatured at 65 °C for 5 min and resolved in 0.5X TBE running buffer at RT. For size standards, we used a 1:1 mix of Low Range ssRNA Ladder (New England Biolabs™, catalog no. N0364S) and 14-30 nt ssRNA Ladder Marker (Takara™, catalog no. 3416). Gels were stained with 1X SYBR Gold Nucleic Acid Gel Stain (Invitrogen™, catalog no. S11494) in 0.5X TBE for 10 min, washed twice with distilled water, and imaged using a Bio-Rad ChemiDoc imaging system.

### DIG-labeled Northern Blots

RNA samples resolved in denaturing polyacrylamide gels were transferred to positively charged nylon membranes (Cytiva, Hybond-N+, catalog no. 45-000-850) using a semi-dry Trans-Blot Transfer System (Bio-Rad, catalog No. 1703940) in 0.5X TBE at constant 20 V for 45 min. Membranes were UV cross-linked twice at 120,000 µJ/cm^2^ for 30 s using a UVC-508 Ultraviolet Cross-linker (Ultra-Lum) and prehybridized for 40 min at 42 °C in DIG Easy Hyb solution (Roche) containing 0.1 mg/mL of Poly(A). Following prehybridization, membranes were hybridized overnight at 42 °C with digoxigenin-labeled DNA probes (2.5 pmol/mL, in DIG Easy Hyb solution + Poly(A)). Oligonucleotides were obtained from Integrated DNA Technologies (IDT, USA), and labeled with DIG Oligonucleotide Tailing Kit, 2nd generation (Roche, catalog no. 03- 353-583-910), following manufacturer’s instructions. After hybridization, membranes were washed twice for 5 minutes at RT with low stringency wash buffer (2X SSC, 0.1% SDS), and twice for 10- 15 min at 42 °C with high stringency wash buffer (1X SSC, 0.1% SDS), blocked for 30-40 min at RT with 1X blocking solution (Roche) in maleic acid buffer (0.1 M maleic acid, 0.15 M NaCl, pH 7.5) and probed for 30 min with an alkaline phosphatase-labeled anti-digoxigenin antibody (Roche, catalog no. 11093274910). Membranes were washed twice for 15 min with washing buffer (0.1 M maleic acid, 0.15 M NaCl, pH 7.5, 0.3% (v/v) Tween 20) and then incubated for 5 min in detection buffer (0.1 M Tris-HCl, 0.1 M NaCl, pH 9.5). Signals were then visualized using CDP-Star ready-to-use (Roche) and detected using the Bio-Rad ChemiDoc imaging system. DNA oligonucleotides used for hybridization probes are listed in *SI Appendix,* Table S1.

### RNA and Protein Quantification

To estimate the concentration of RNA isolated from cell lysates and AWFs, a NanoDrop (Thermo Fisher Scientific™) instrument was used. For LSW RNA, we estimated RNA concentrations using densitometric quantification of SYBR Gold-stained polyacrylamide gels. This was necessary because LSW samples contained a contaminant that caused an overestimation of RNA concentration when using absorbance-based quantifications. Briefly, 1 µL of RNA from LSW and AWF was loaded in a denaturing urea-polyacrylamide gel and run as previously described. The gel was then stained with SYBR Gold stain and imaged using the Bio-Rad ChemiDoc imaging system. The densitometric ratio between AWF and LSW was calculated using ImageJ software (National Institutes of Health, USA) (55), and this ratio along with the nanodrop value of AWF RNA was used to calculate the concentration in ng/µL of the LSW RNA.

To determine protein concentrations in AWF, we employed the Bradford method (56) using bovine serum albumin as a standard. Due to the extremely low concentration of proteins in LSW samples, we had to perform densitometric quantification upon silver-staining of denaturing polyacrylamide gels. Briefly, an equivalent volume of LSW and AWF per gram FW from each replicate was loaded in an SDS-PAGE gel (12% w/v acrylamide) and run at 100 V for 1 h. Gels were then stained with the ultra-sensitive Pierce Silver Stain Kit (Thermo Fisher Scientific™, catalog no. 24612) following the manufactureŕs instructions. Gel images were acquired using the Bio-Rad ChemiDoc Imaging System. The densitometric ratio between AWF and LSW was calculated using the ImageJ software, and this ratio and Bradford estimation of AWF proteins were used to estimate the protein concentration of LSW samples.

### Immunoblots

Proteins from equivalent volumes of AWF and LSW per gram FW were precipitated by mixing with 4 volumes of ice-cold acetone and incubating on ice for at least 1 h or overnight at –20 °C. Then, the samples were centrifuged at 14,000 *g* for 10 min at 42 °C. The supernatant was discarded, and then the tubes were inverted to allow the residual acetone to evaporate from the pellet. The pellet was allowed to dry for no longer than 1 h and resuspended in filtered VIB.

For immunoblots, 30 µL of resuspended pellets were combined with 10 µL of 4X SDS loading buffer (250-mM Tris–HCl, pH 6.8, 8% (w/v) sodium dodecyl sulfate, 40% (v/v) glycerol, 20% 2-mercaptoethanol and 0.004% (w/v) bromophenol blue) and were then denatured at 95 °C for 5 min. Then, 40 µL of each sample was loaded onto stain-free gradient gels (4–20% Precise Protein Gels, Thermo Scientific) and separated at 150 V for 1 h in 1X TBS electrophoresis running buffer (24.8 mM Tris base, 0.1% (w/v) sodium dodecyl sulfate, 192 mM glycine, pH 8.3). The resolved proteins were visualized using the Bio-Rad ChemiDoc imaging system. After the proteins were transferred to a nitrocellulose membrane (Amersham™ Protran® Premium Western blotting membrane, nitrocellulose) using the semi-dry Trans-Blot Transfer System in Transfer buffer, the membrane was stained with Ponceau S stain (0.1% dye in 5% acetic acid solution) for about 10 min and cleared with water to visualize the protein transfer.

The membrane was washed once with 1X Tris-buffered saline (50 mM Tris-Cl and 150 mM NaCl, pH 7.5) containing 0.1% Tween-20 (TBST) before blocking with 5% (w/v) Difco Skim Milk (BD) prepared in 1X TBST for 1.5 h at RT. Thereafter, the membrane was incubated overnight at 4 °C with the following primary antibodies at the indicated dilutions: rabbit polyclonal anti-PATL1 ((57); 1:5,000), rabbit polyclonal anti-ESM1, rabbit polyclonal anti-ANN2, rabbit polyclonal anti- ANN1, rabbit polyclonal anti-PEN1 ( (58); 1:1,000), rabbit polyclonal anti-PR5, mouse monoclonal [9F9.F9] anti-GFP (Abcam, Cambridge, UK, catalog no. ab1218; 1:2,000). Membranes were then washed with 1X TBST, and if needed, incubated with one of the following secondary antibodies as appropriate: Horseradish peroxidase (HRP)-conjugated goat anti-rabbit (Abcam, catalog no. ab97051; 1:10,000) or HRP-conjugated goat anti-mouse (Abcam, catalog number ab6789; 1:5,000) for 1.5 h at RT. After three final washes in 1X TBST, proteins were visualized using ProtoGlow ECL Substrate (National Diagnostics) and the Bio-Rad ChemiDoc Imaging System. The densitometric ratio between AWF and LSW was calculated using the ImageJ software.

### RNase A and RNase R Treatment of RNA samples

RNA was isolated from AWF using TRIzol as described previously and resuspended in ultrapure water. For RNase R treatment, 200 ng of RNA were treated with 3 units of RNase R (Lucigen. RNR07250) in a 10 uL reaction volume containing 1X RNaseR reaction buffer for 1 h at 37 °C. RNase R was then inactivated by incubation at 65 °C for 20 minutes. For RNase A treatment, 200 ng of total RNA were treated with 20 ng/µL of RNase A (Thermo Scientific, EN0531) diluted in 1x reaction buffer (15 mM NaCl, 10 mM Tris-HCl, pH 7.5) in a 10 µL reaction volume for 1 hour at room temperature. RNase A activity was stopped by adding a mixture of RNase Inhibitor, Murine (APExBIO) and RNase Out (Invitrogen). Following RNase treatments, RNA was purified by precipitation with ammonium acetate and ethanol to remove free nucleotides and other small degradation products. This precipitation step was repeated twice for optimal RNA purity.

### Quantification of N^6^-Methyladenosine (m^6^A) in Extracellular RNA

RNA was isolated from LSW and AWF using TRIzol as described above, and the RNA concentrations were measured using a ThermoFisher NanoDrop One spectrophotometer. For all samples, equal amounts of RNA were prepared in equal volumes (6 µL) using UltraPure DNase/RNase-free distilled water (Invitrogen). RNA samples were denatured at 95 °C for 3 min and placed on ice immediately to prevent the formation of secondary structures. m^6^A quantification using dot blots was performed using the protocol described by Zand Karimi *et al.* (11). Briefly, RNA samples were applied directly to a piece of Hybond-N+ membrane (Amersham Pharmacia Biotech) using a micropipette. To prevent the spread of RNA on the membrane, 2 µL of RNA solution was applied at a time, allowing the membrane to dry for three min before applying the next 2 µL drop to the same spot until a total of 6 µL of RNA sample was applied. To crosslink the spotted RNAs to the membrane, a UVC-508 Ultraviolet Cross-linker (Ultra-Lum) was used to irradiate the membrane twice at 120,000 microjoules/cm2 for 30 s. The membrane was then washed in clean RNase-free 1X PBS buffer (1xPBS; 2.7 mM KCl, 8 mM Na2HPO4, 2 mM KH2PO4, and 137 mM NaCl, pH 7.4) and blocked in 5% (w/v) non-fat milk prepared in 1X PBS containing 0.02% Tween-20 for 1 h at RT. The membrane was then incubated overnight with anti-m^6^A antibody (Abcam, catalog no. Ab151230; or Synaptic Systems, catalog no. 202 003) at a 1:250 dilution in 5% non-fat milk prepared in 1X PBS containing 0.02% Tween-20. The membrane was washed in 1X PBS containing 0.02% Tween-20 three times and incubated with horseradish peroxidase-labeled goat anti-rabbit antibody (Abcam, catalog no. ab205718) at a 1:5,000 dilution for 1 h at RT. After a final wash in 1X PBS containing 0.02% Tween- 20, m^6^A modified RNAs were visualized using the Immune-Star Reagent (Bio-Rad) and imaged using the Bio-Rad ChemiDoc Imaging System.

Alternatively, EpiQuik™ m^6^A RNA Methylation Quantification Kit (Fluorometric) (Catalog no. P-9008-48, EpigenTek) was used to determine the m^6^A percentage in exRNA by following the instructions provided in the user guide. Briefly, total RNA is bound to strip wells using an RNA high-binding solution. m^6^A is detected using a specific capture N6-methyladenosine (anti-m6A) antibody and detection antibody. The detected signal is enhanced and then quantified colorimetrically by reading the absorbance in a microplate spectrophotometer at a wavelength of 450 nm. The amount of m^6^A is proportional to the OD intensity measured. Both negative and positive RNA controls provided in this kit must be used to quantity the percentage of m^6^A. This kit allowed us to quantify the absolute amount of m^6^A in each sample.

### Detection of Pectin in Extracellular RNA Fractions

RNA was isolated from LSW and AWF using TRIzol as described above. For all samples, 100 ng of RNA were prepared in equal volumes (4 µL) using UltraPure DNase/RNase-free distilled water (Invitrogen). RNA samples were denatured at 95 °C for 3 min and placed on ice immediately to prevent the formation of secondary structures. Dot blot analysis of these RNA samples was performed using the same protocol as for m6A quantification except using JIM7 as the primary antibody at a dilution of 1:10 (Kerafast, catalog no. ELD005) and horseradish peroxidase-labeled goat anti-rat as the secondary antibody at a dilution of 1:5000 (Invitrogen, catalog no. 31470).

### Statistical Analyses

Statistical analyses and plotting of RNA and protein concentrations were performed using the GraphPad Prism 8.3.0 software (GraphPad Software, San Diego, CA, USA). The specific statistical test used for each analysis is provided in the corresponding figure legend. The number of independent biological replicates (n) is indicated in each plot.

### Preparation of sRNA Sequence and Standard RNA Sequence Libraries

sRNA libraries were constructed using the RealSeq-AC kit version 2 (Realseq Biosciences, Santa Cruz, CA, USA, catalog no. 500-00048;) as per the manufacturer’s instructions. We used 60 ng of DNase I-treated total RNA as starting material for constructing libraries. (DNaseI: catalog no. EN0521; Thermoscientific). For the Cell lysate rRNA depleted samples, we used the RiboMinus Plant Kit for RNA-seq (Invitrogen, catalog no. A10838-08) at 1/10^th^ of the recommended volume of regents and sample. For RNAseq libraries, we used NEBNext Ultra II Directional RNA Library Prep Kit for Illumina (New England Biolabs, catalog no. E7760L) protocol. To ensure correct library size capture, we performed a Bioanalyzer dsDNA HS chip assay on the Agilent 2100 Bioanalyzer (no. DE4103649, Agilent Technologies, Inc.) for each library. We sent all sRNA and RNAseq libraries for sequencing to the University of Delaware Sequencing and Genotyping Center, where they were sequenced on a NovaSeq2000 instrument using 50-bp single-end reads for sRNA and 75-bp paired-end reads for RNAseq.

### Sequence Data Analysis

All sRNA libraries were analyzed as previously (11). Briefly, we first trimmed the adaptors using Cutadapt version 1.16 (59) using a minimum insert size of 10 nt and no maximum. We assessed sequence quality using FastQC (http://www.bioinformatics.babraham.ac.uk/projects/ fastqc/). We aligned clean reads to the Arabidopsis genome (TAIR version 10) and all subsequent analyses were performed using the software Bowtie2 (60). For miRNA analyses, we used the latest version of miRBase (version 22; (61)). Sequences in the RNA-seq libraries were analyzed using HiSat2 and Stringtie pipeline (43), and the TAIR10 available annotation file. We performed differential accumulation analyses using DESeq2 with default parameters, using reads that were not normalized as input (62). In DESeq2, p-values were calculated using the Wald test and corrected for multiple testing using the Benjamini and Hochberg procedure (63). We generated graphical representations using the software ggplot2 (64) in the R statistical environment.

To analyze the tRNA derived sRNAs from the sRNAseq data, we ran unitas version 1.8.0 (36) using the Genomic tRNA database for Arabidopsis (data accessed on 05 March 2024) (65, 66) using default parameters. The absolute read counts from the unitas pipeline were normalized against total number of input reads per million. The plots were drawn using R.

## ACKNOWLEDGMENTS

We thank Dorothee Staiger at the University of Bielefeld for providing seed of GRP7-GFP transgenic Arabidopsis and for supplying anti-GRP7 antisera, the Indiana University Physical Biochemistry Instrumentation Facility for access to ultracentrifuges and nanoparticle tracking equipment, and the University of Delaware Sequencing and Genotyping Center for assistance with the generation of sRNA-seq and RNA-seq data. We also thank David Daleke and Craig Pikaard for providing access to centrifuges, and members of the Innes and Meyers laboratories for valuable discussions. This work was supported by three grants from the United States National Science Foundation, IOS-2243531 and IOS-1842685 to RWI and IOS-1842698 to BCM and PB, and IOS- 2243534 to PB.

## Author Contributions

L.B., M.S.-R., and R.W.I. designed the research; L.B., M.S.-R., P.B., H.Z.K. and M.M. performed the research; P.B. and M.S.-H analyzed sequence data; and L.B., M.S.-R. and P.B. wrote the paper with editing by B.C.M. and R.W.I.

## Supplementary Figures and Tables

**Fig S1.**
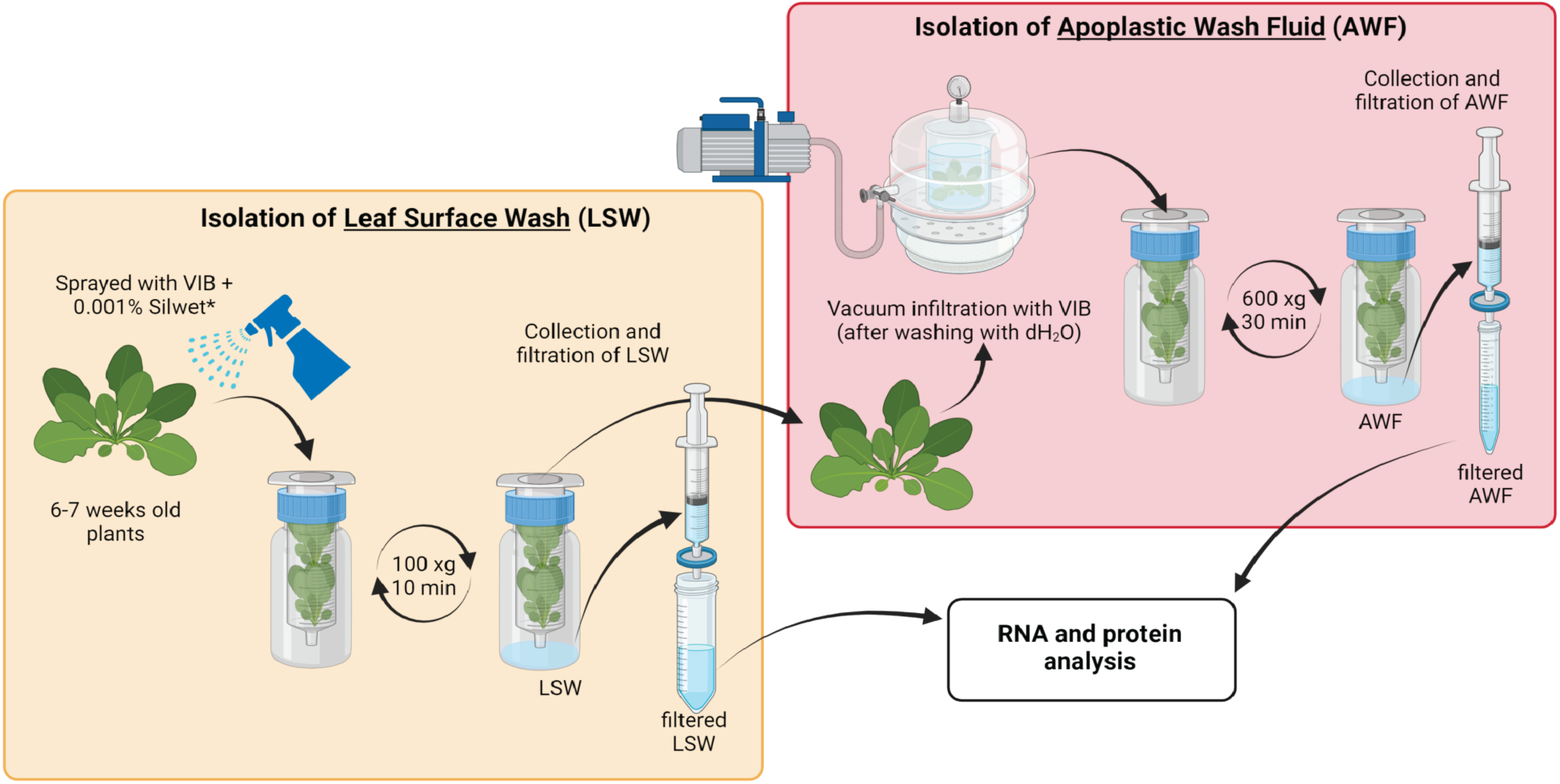
Schematic illustration of the stepwise protocol for the isolation of LSW and AWF using Arabidopsis plants. Full rosettes of six-to-seven-week-old Col-0 plants were detached and sprayed on both sides with VIB supplemented with 0.001% (v/v) Silwet. Rosettes were then placed in a 60 mL syringe with small holes that was inserted into a 250 mL centrifuge bottle. The rosettes were then centrifuged at 100 *g* for 10 mins at 4 °C. To isolate the AWF, the same set of plants was then vacuum infiltrated with VIB followed by centrifugation at 600 *g* for 30 mins at 4 °C. Both fractions were filtered through 0.2 µm filters before further processing. The weight of the rosettes was also recorded to normalize the concentration of RNA and protein isolated from both AWF and LSW based on plant fresh weight (FW).

**Fig S2.**
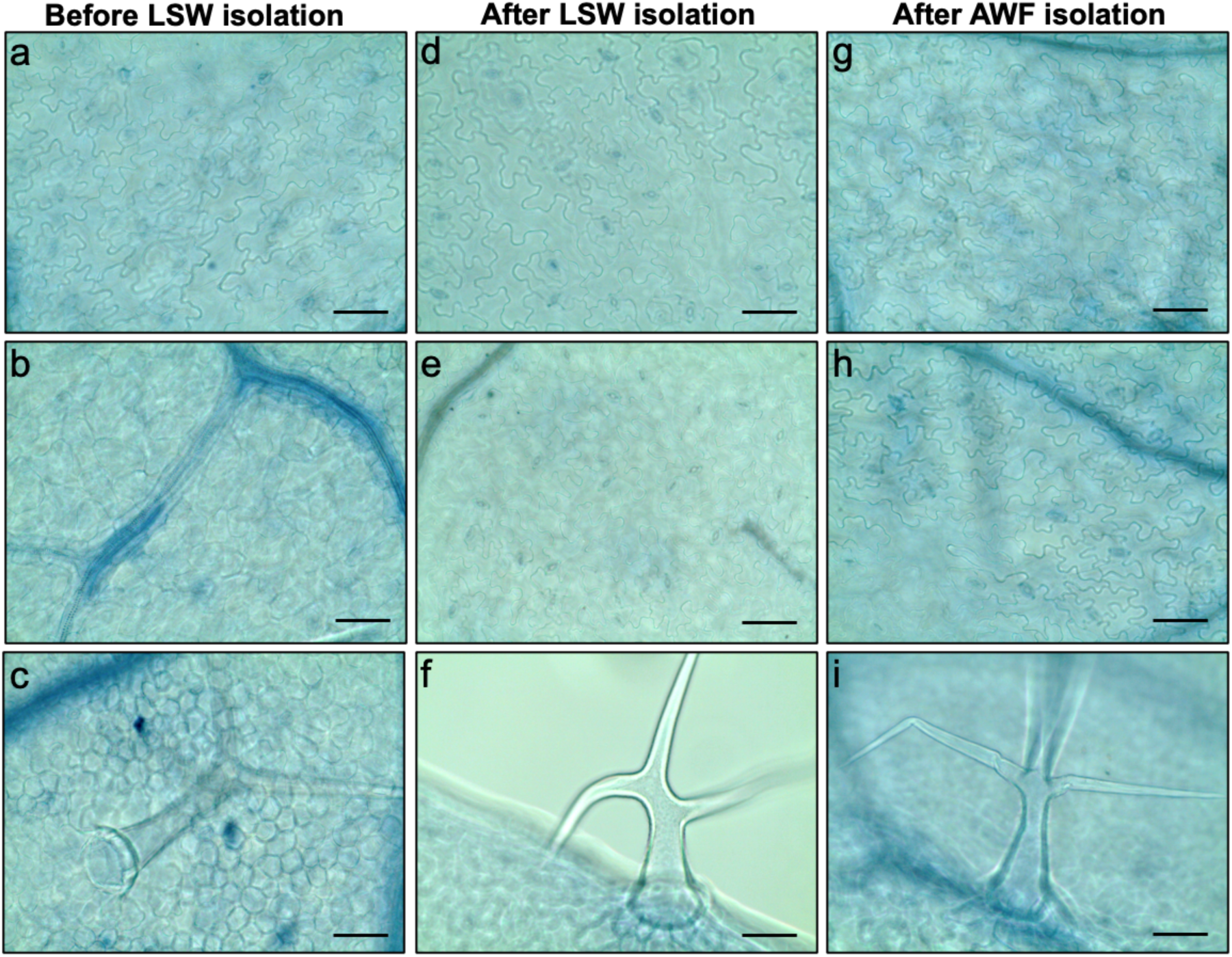
Isolation of LSW and AWF does not cause cell rupture. To assess if LSW and AWF isolation leads to cell rupture, leaves from three whole rosettes were stained with trypan blue dye before and after each isolation step as indicated. *(a-c)* Before LSW isolation; *(d-f)* After LSW isolation; *(g-i)* After AWF isolation.

**Fig. S3.**
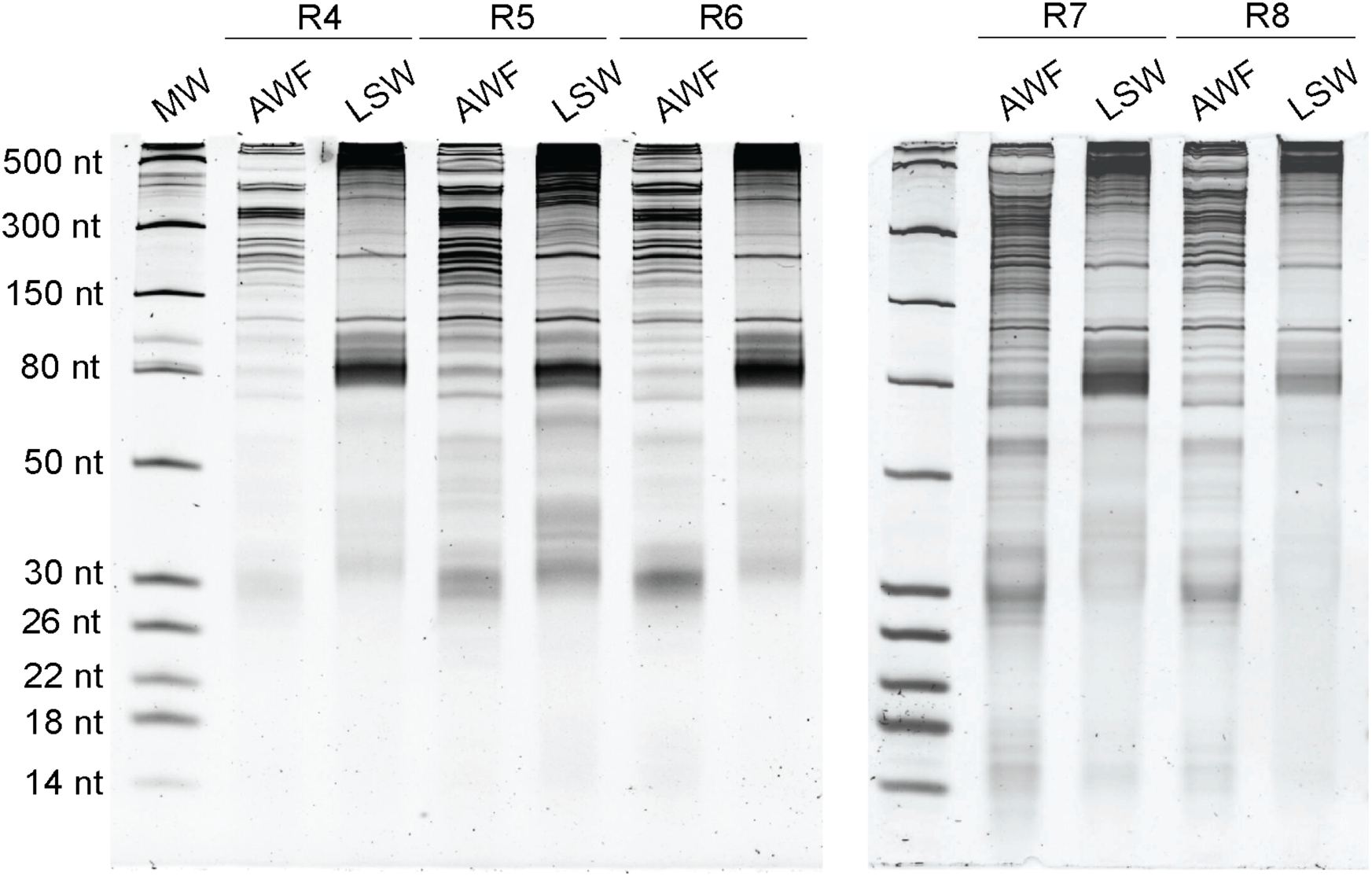
AWF and LSW contain equivalent amounts of RNA. RNA isolated from AWF and LSW from eight different replicates (R1-R3: shown in Fig. 1*C*) was separated on 15% denaturing polyacrylamide gels and stained with SYBR GOLD nucleic acid stain. Densitometry analysis was performed using ImageJ software to estimate the amount of RNA per gram fresh weight of plant material (Refer to Fig. 1*C* for the plot of this analysis).

**Fig. S4.**
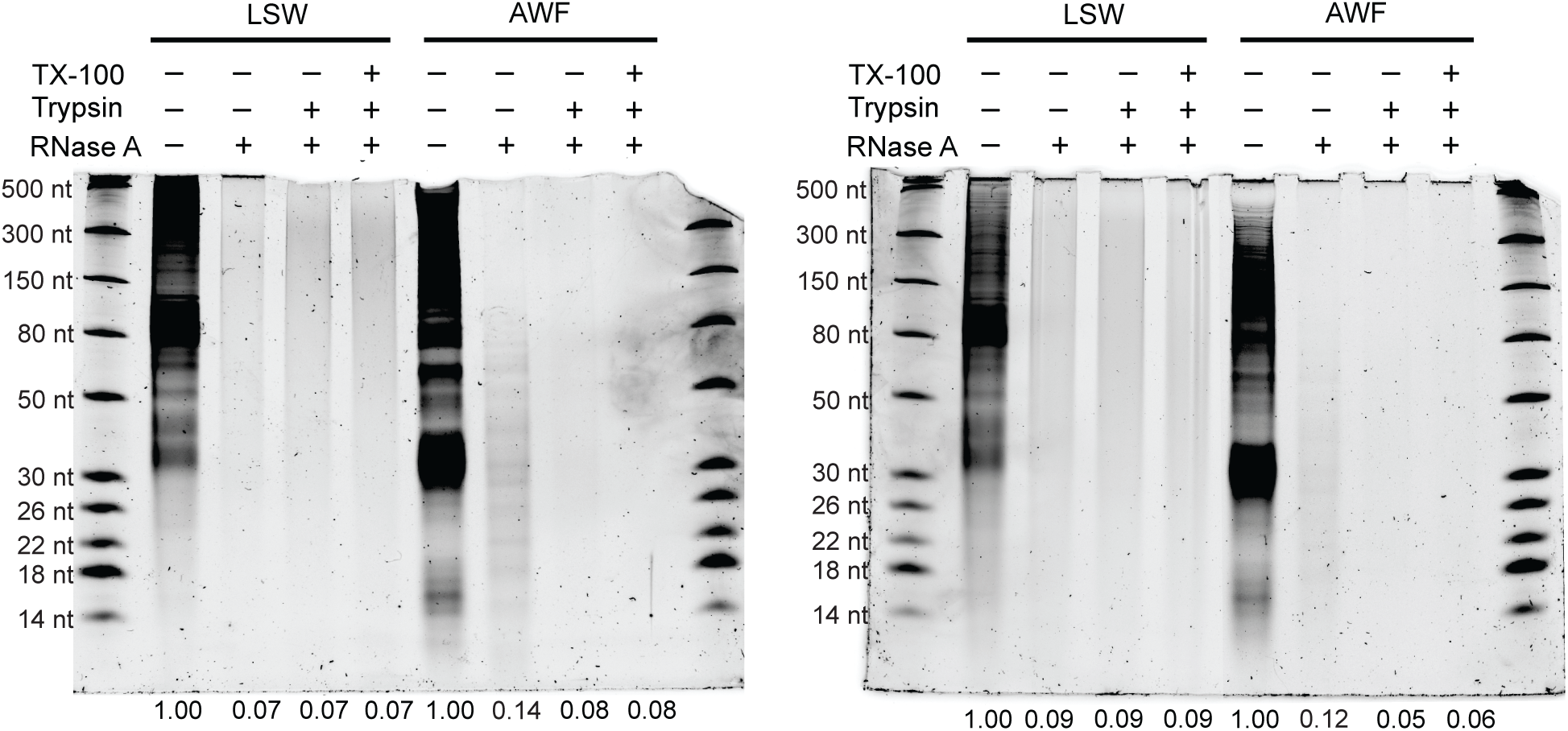
Ribonuclease protection assays of AWF and LSW. (Replicates 2 and 3 of the experiment presented in Fig. 2*A*). AWF and LSW samples were treated with RNase A, or trypsin followed by RNase A, or TX-100 followed by trypsin followed by RNase A at RT. The negative control was mock-treated and placed on ice. RNAs were extracted using TRIzol, separated on a 15% denaturing polyacrylamide gel, and stained with SYBR Gold nucleic acid stain. Numbers along the bottom of gel image indicate relative RNA abundance in each lane, which was estimated by densitometry and expressed relative to the mock-treated sample.

**Fig. S5.**
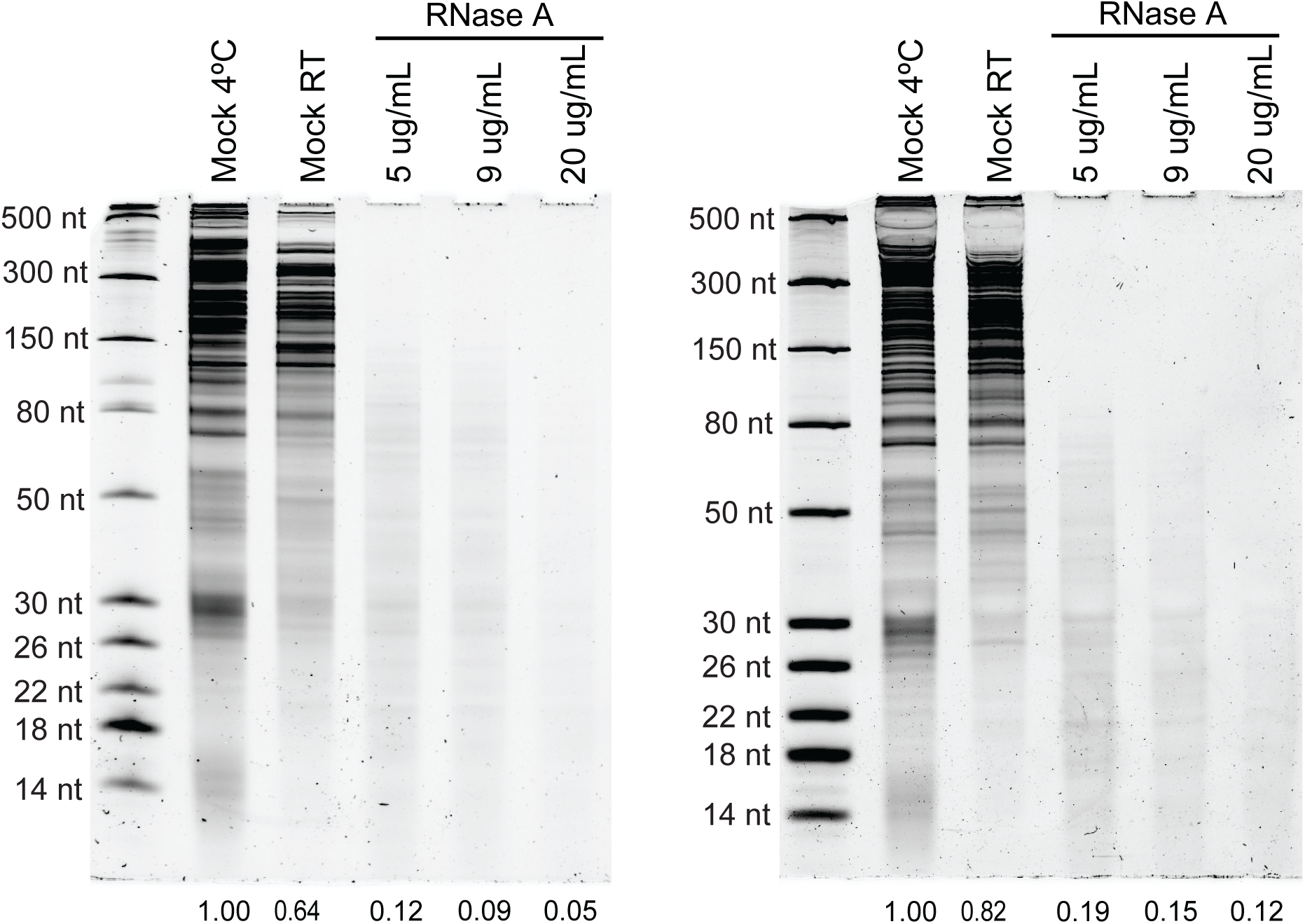
RNA molecules in AWF are partially protected from endoribonuclease digestion by proteins. AWF samples were treated with increasing concentrations of RNase A as indicated. Treatments were performed at RT for 1 h. Mocks were treated with the buffer used to dilute RNase A, subjected to the same incubation time as RNase A treated samples, and kept either at RT or on ice (4 °C). RNAs were extracted using TRIzol, separated on a 15% denaturing polyacrylamide gel, and stained with SYBR Gold nucleic acid stain. RNA abundance in each gel lane was estimated by densitometry and expressed relative to Mock (4 °C). This experiment was repeated at least three times with similar results (two biological replicates are presented). Biological replicates were isolated on different days.

**Fig. S6.**
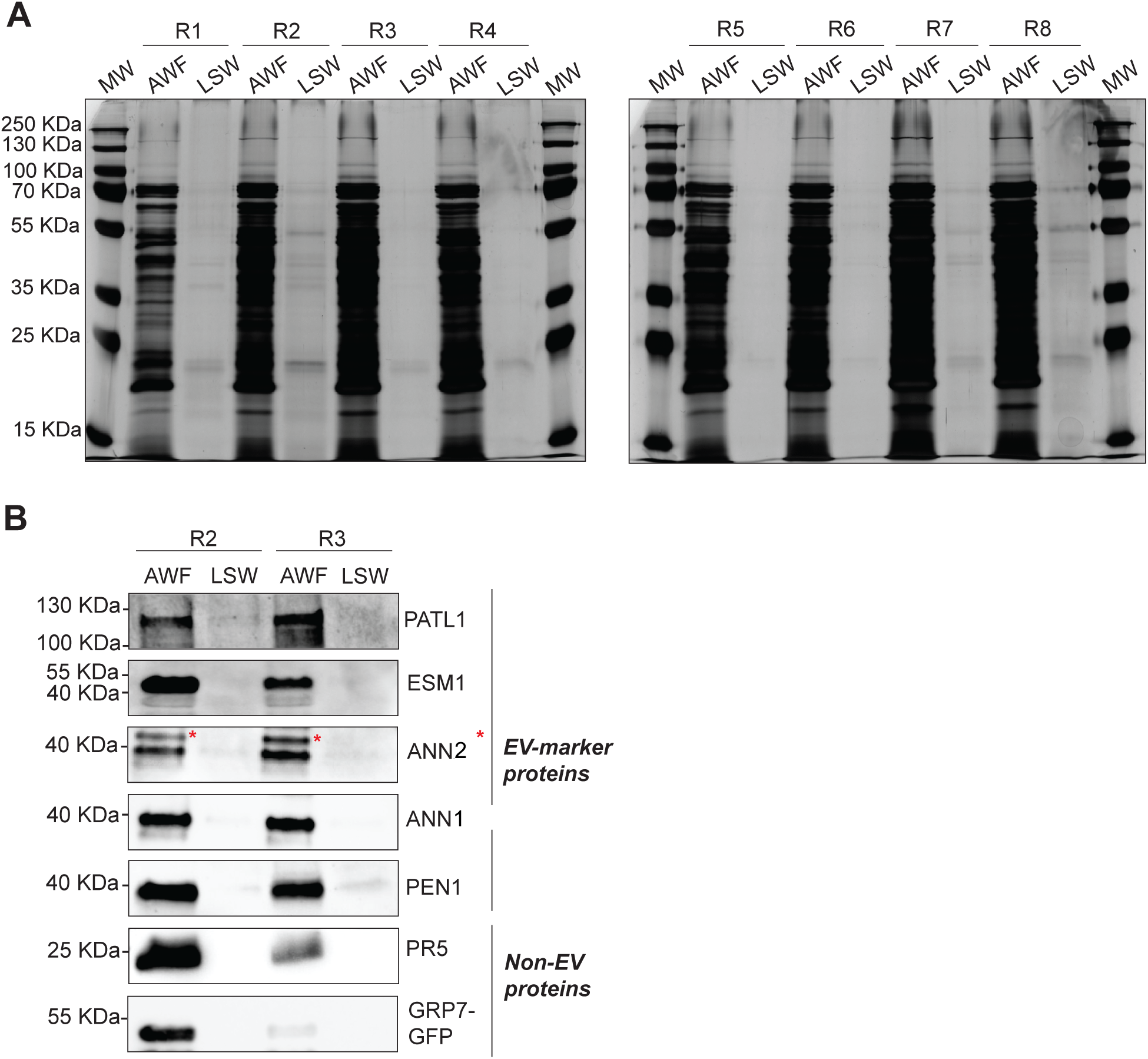
LSW contains very little protein compared to AWF. (*A*) Silver-stained SDS-PAGE showing the total protein profile of AWF and LSW. Eight biological replicates were assayed to estimate the amount of protein per fresh weight in both fractions (refer to Fig. 1*B* for this analysis). The amount of protein loaded in each lane was normalized by leaf fresh weight. MW: molecular weight markers. (*B*) Immunodetection of EV-marker proteins (PEN1, PATL1, ESM1, ANN1, ANN2) and RNA-binding proteins (GRP7, PR5, ANN1, ANN2) in AWF and LSW. These are biological replicates 2 and 3 of the experiment presented in Fig. 2*C*. Red asterisks indicate the specific band for ANN2.

**Fig. S7.**
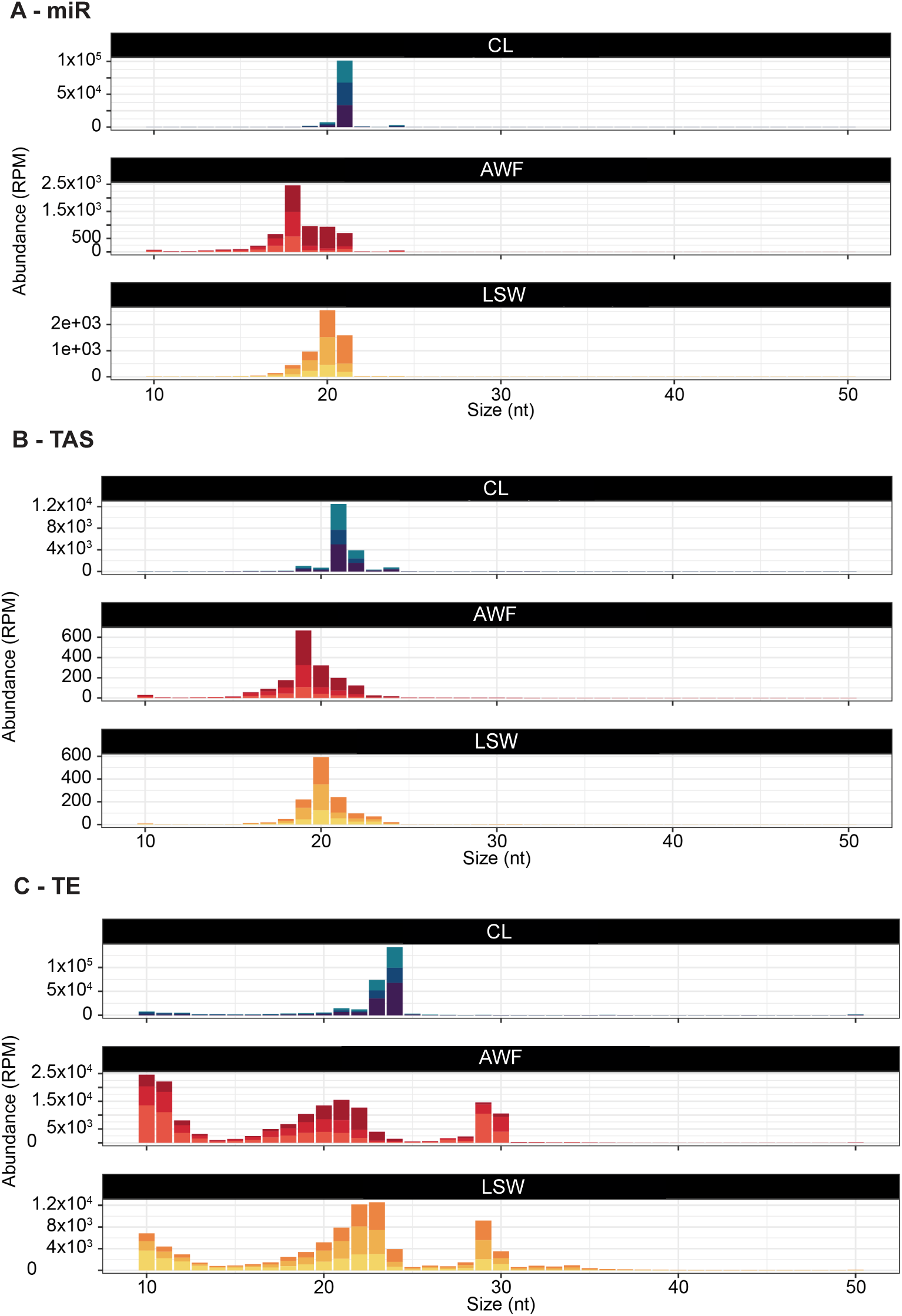
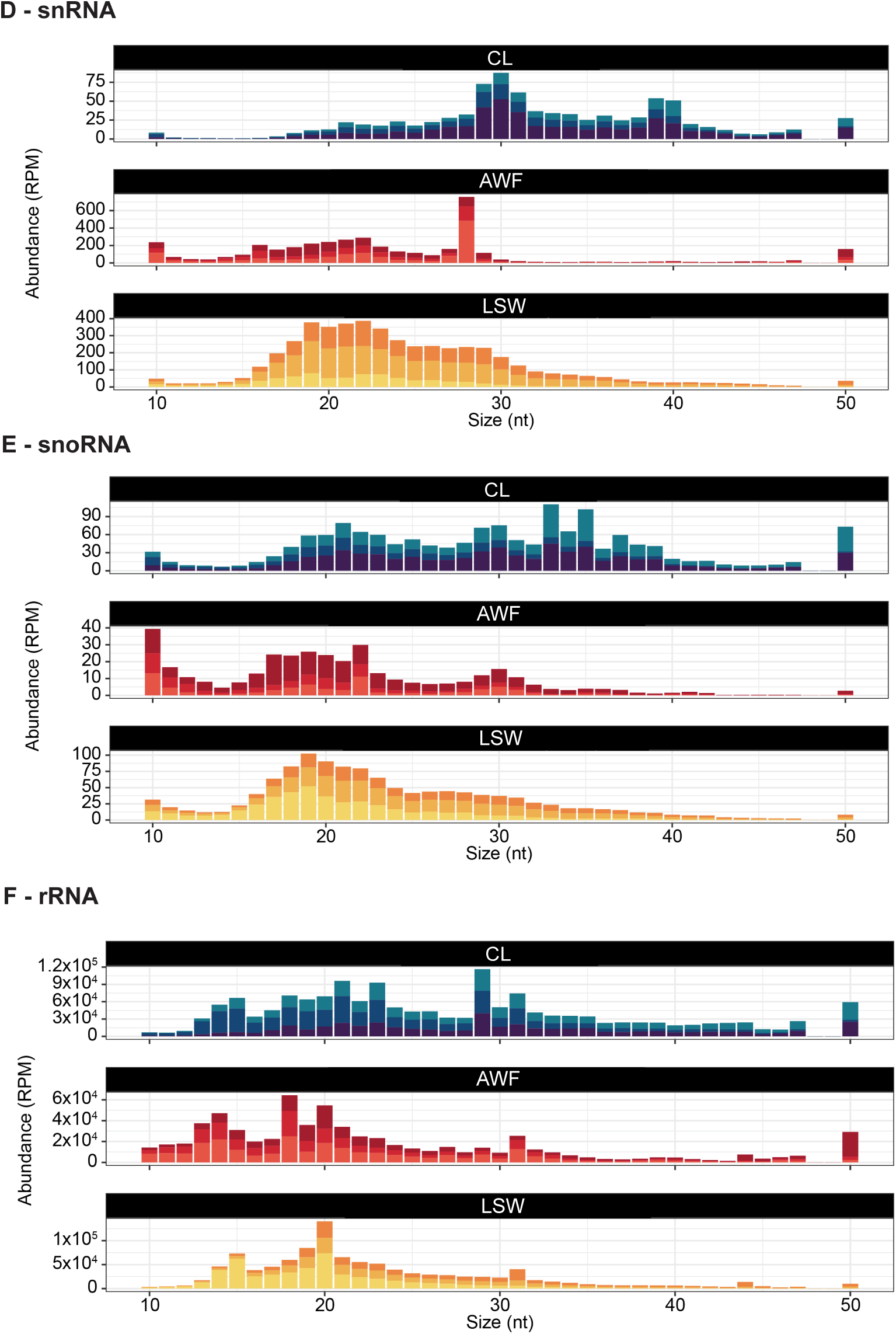
Size distribution of sRNAs categorized by source. sRNA size distribution of reads mapping to the Arabidopsis genome (TAIR version 10). The abundance of each size class was calculated for each sample independently and normalized to the total number of reads for that sample. The *x*-axis indicates the sRNA size, from 10 to 50 nucleotides, and the *y*-axis indicates its abundance in reads per million (RPM) reads. Each panel represents an RNA fraction, from top to bottom, CL, AWF, and LSW. (*A*) Reads mapping to microRNAs (miR). (*B*) Reads mapping to trans-acting small interfering RNAs (tasiRNAs). (*C*) Reads mapping to transposable elements (TE). (*D*) Reads mapping to small nuclear RNAs (snRNAs). (*E*) Reads mapping to small nucleolar RNAs (snoRNAs). (*F*) Reads mapping to ribosomal RNAs (rRNAs).

**Fig. S8.**
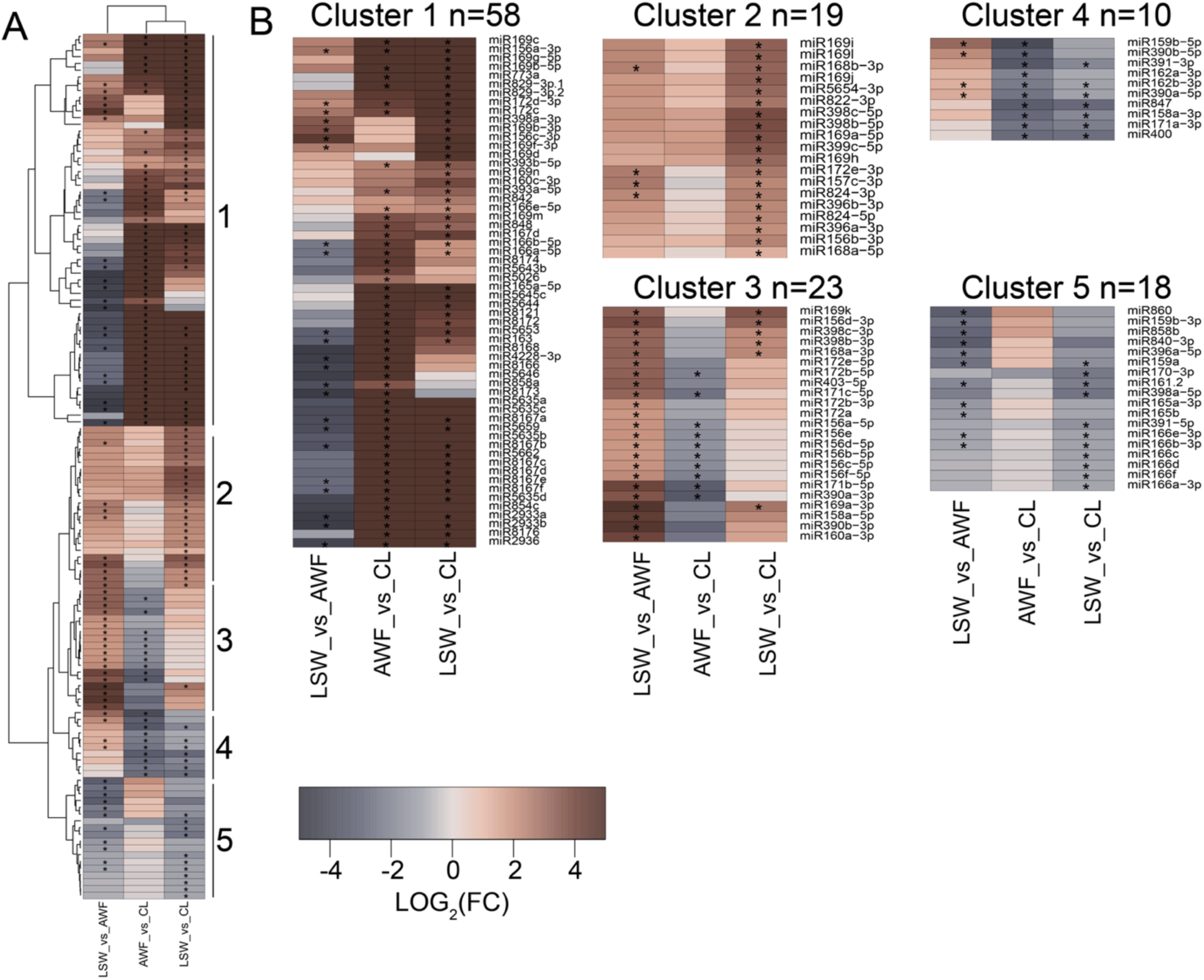
Specific miRNAs differentially accumulate in LSW and AWF. The heatmap representing all differentially accumulated microRNAs is divided into three columns with enrichment indicated in brown shades and depletion indicated in grey shades. The first column indicates LSW compared to AWF, the second column indicates AWF compared to CL and the third column indicates LSW compared to CL. (*A*) Includes all differentially accumulating miRNAs. (*B*) Details of each of the five clusters.

**Fig. S9.**
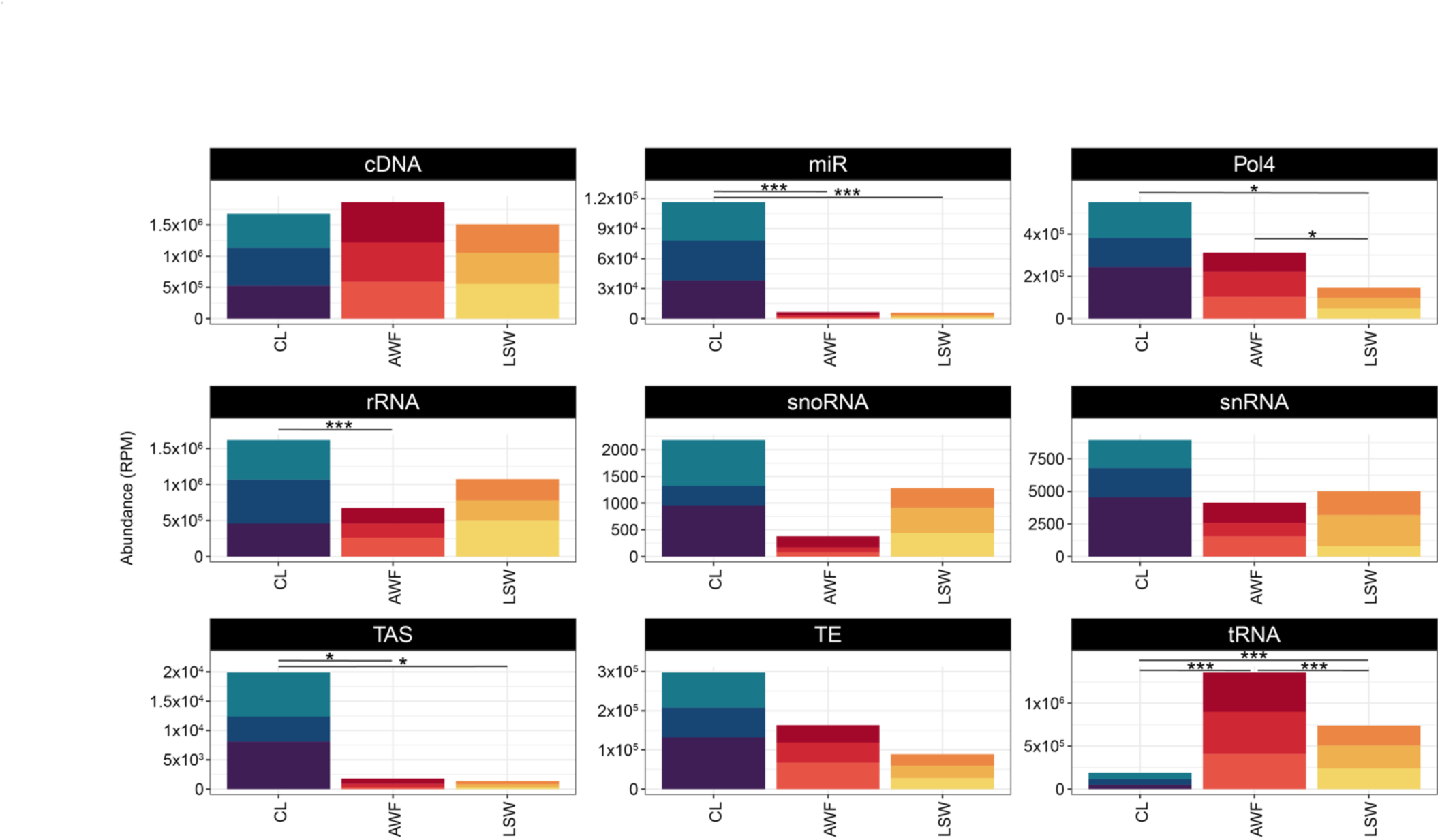
Sources of sRNA reads obtained from CL, AWF, and LSW. Genomic origin of small RNA reads based on the categories established in the TAIR 10 genome version. RNAs that mapped to the genome were categorized by origin. The x-axis represents each of the fractions, CL, AWF, and LSW. The y-axis indicates the relative abundance expressed in RPM. Each box represents a specific genomic source: cDNA, complementary DNA; Pol4, Polymerase IV-dependent products; rRNA, ribosomal RNAs; TE, Transposable elements; tRNA, transfer RNAs; miR, microRNAs; snoRNA, small nucleolar RNA; snRNA, small nuclear RNAs; TAS, trans-acting siRNA. The three colored shades represent three independent biological replicates, with each replicate derived from 18 Arabidopsis plants.

**Fig. S10.**
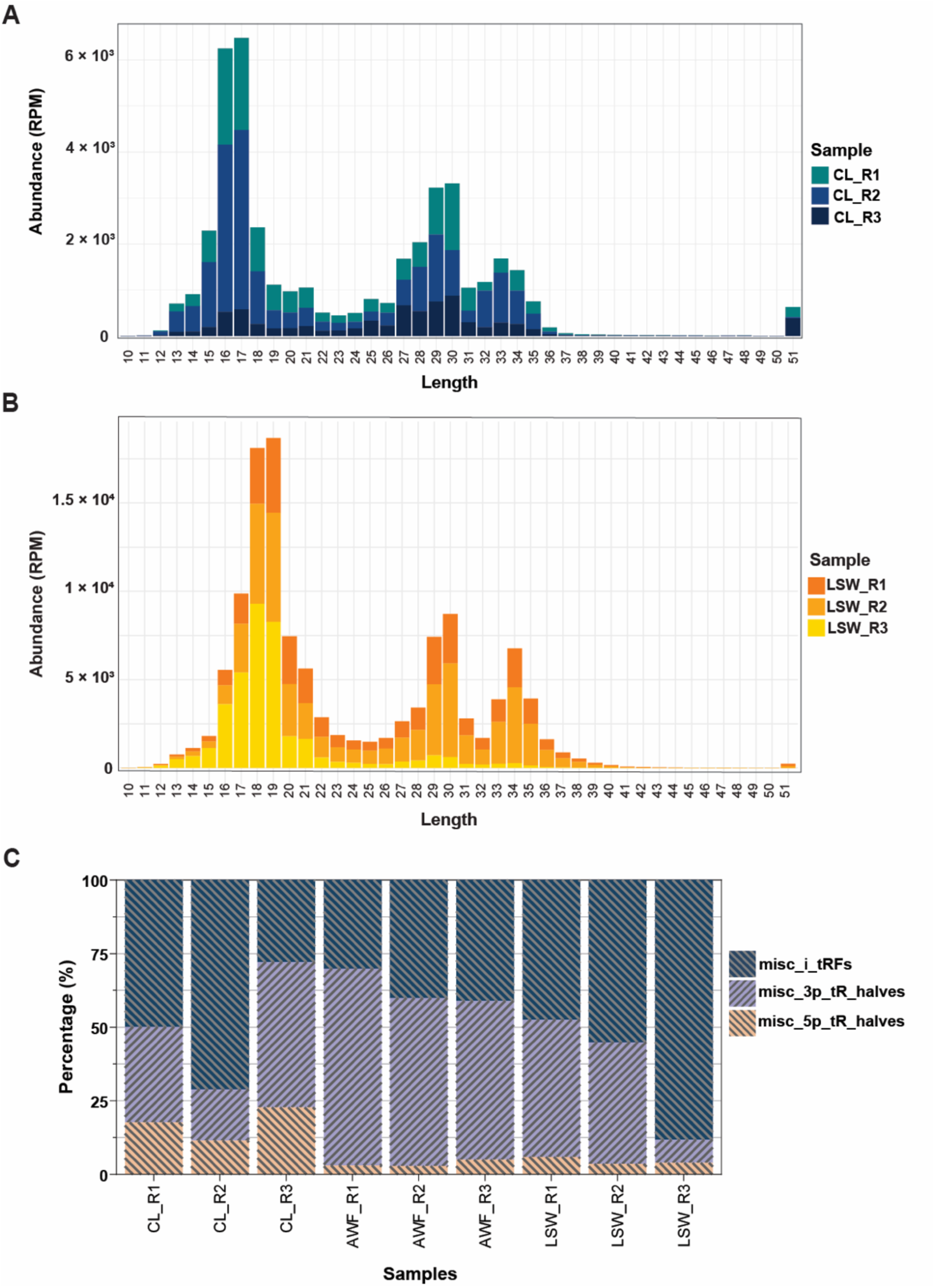
misc-tRFs are enriched in tRNA halves. (*A*) Size distribution of misc-tRF reads in CL samples. The colors represent three replicates. (*B*) Size distribution of misc-tRF reads in LSW samples. The colors represent three replicates. (*C*) Reclassification of the misc-tRFs. misc_5p_tR_halves: start position at nucleotide position 2, 3, or 4 and length 28-35 nt; misc_3p_halves: start position after nucleotide position 29 and length ≥28 nt; and misc_i_tRFs: all misc_tRFs that were not reclassified as either misc_5p_tR_halves or misc_3p_tR_ halves.

**Fig. S11.**
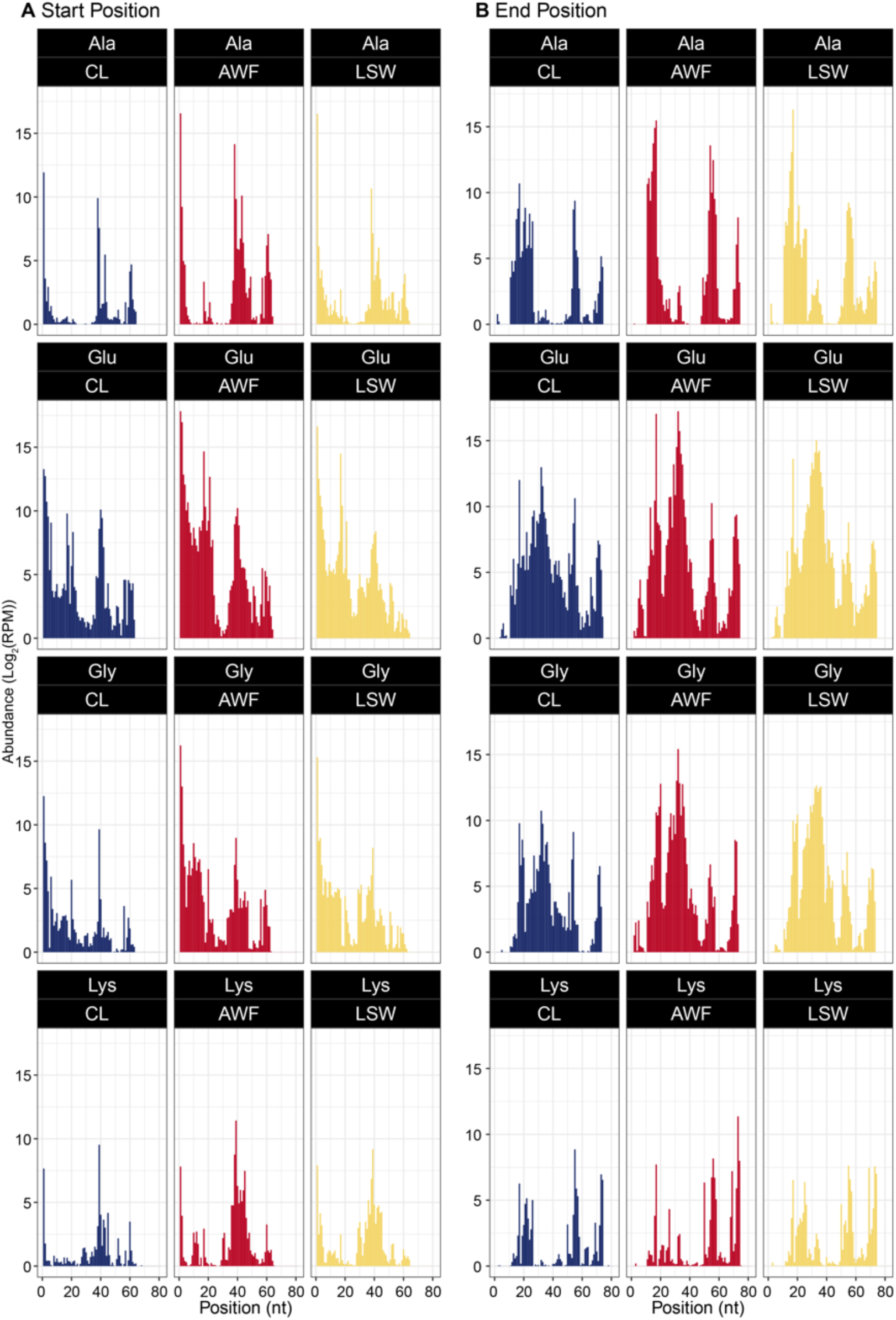
5’ and 3’ ends of tRNA-derived fragments show correlation with Northern blot analysis. The starting and ending positions of small RNA reads mapping to each of the indicated tRNAs is represented using their 5’ end (*A*) and 3’ end (*B*) positions relative to the full-length tRNA. The x-axis represents the position along the tRNA transcript, using 1 as the 5’ end. The y-axis represents the cumulative abundance in logarithmic scale in base 2 of reads per million (RPM). For each plot, the three panels represent the three fractions, CL, AWF, and LSW.

**Fig. S12.**
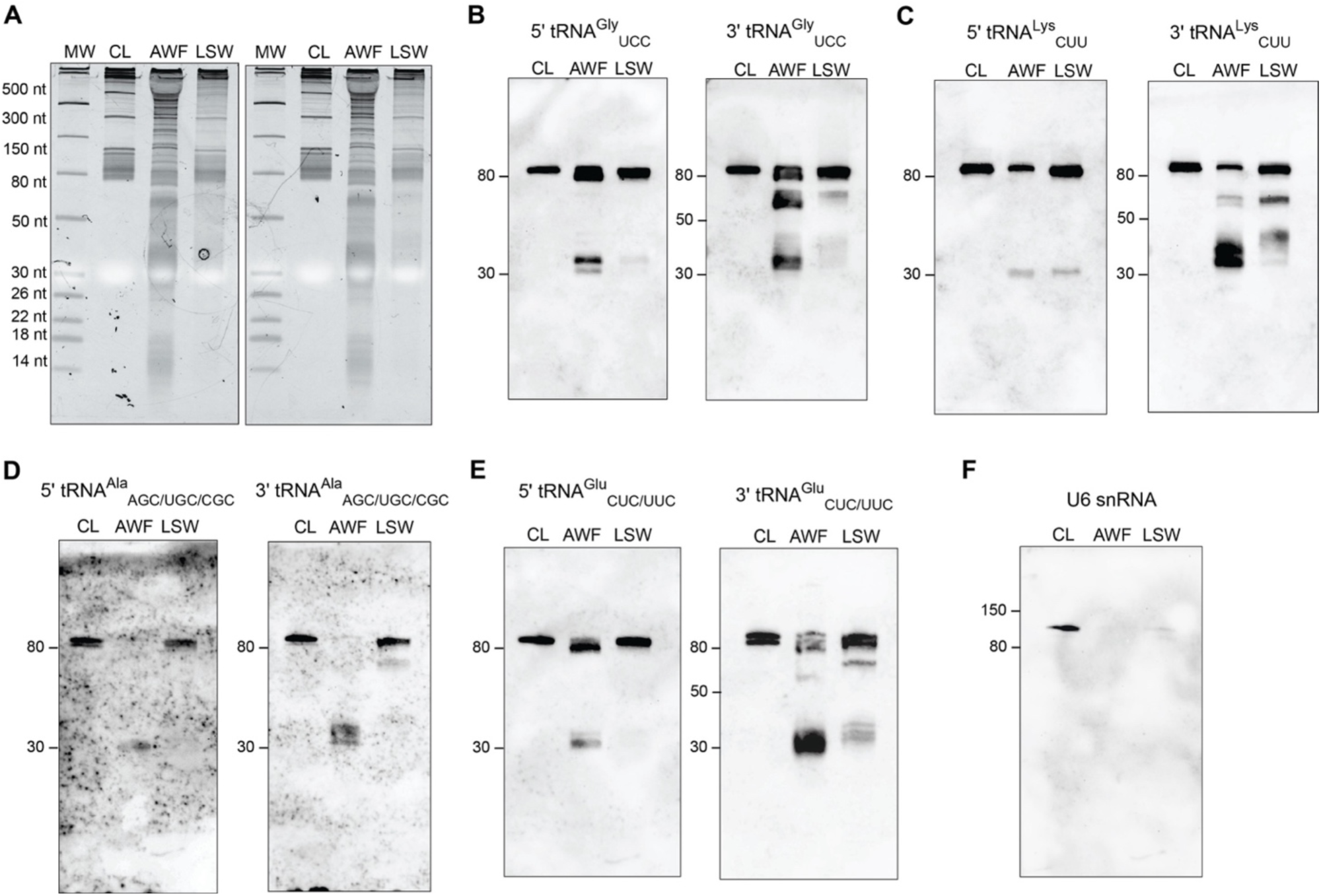
Extracellular fractions are enriched in tRNA-derived fragments compared to whole cell lysate. (*A*) 100 ng of RNA from total CL, AWF, and LSW was separated on 15% denaturing polyacrylamide gels and stained with SYBR GOLD nucleic acid stain. (*B, C, D,* and *E*) Upon blotting onto a positively charged nylon membrane, RNA was probed with DIG-labeled 5’ and 3’ probes against tRNA^Gly^, tRNA^Lys^, tRNA^Ala^, and tRNA^Glu^ to detect tRNA-derived fragments. Full-length tRNAs were also detected in these samples. (*F*) RNA was probed with a DIG-labeled probe against U6 snRNA as a control. This figure shows biological replicates distinct from those shown in Fig. 6.

**Fig. S13.**
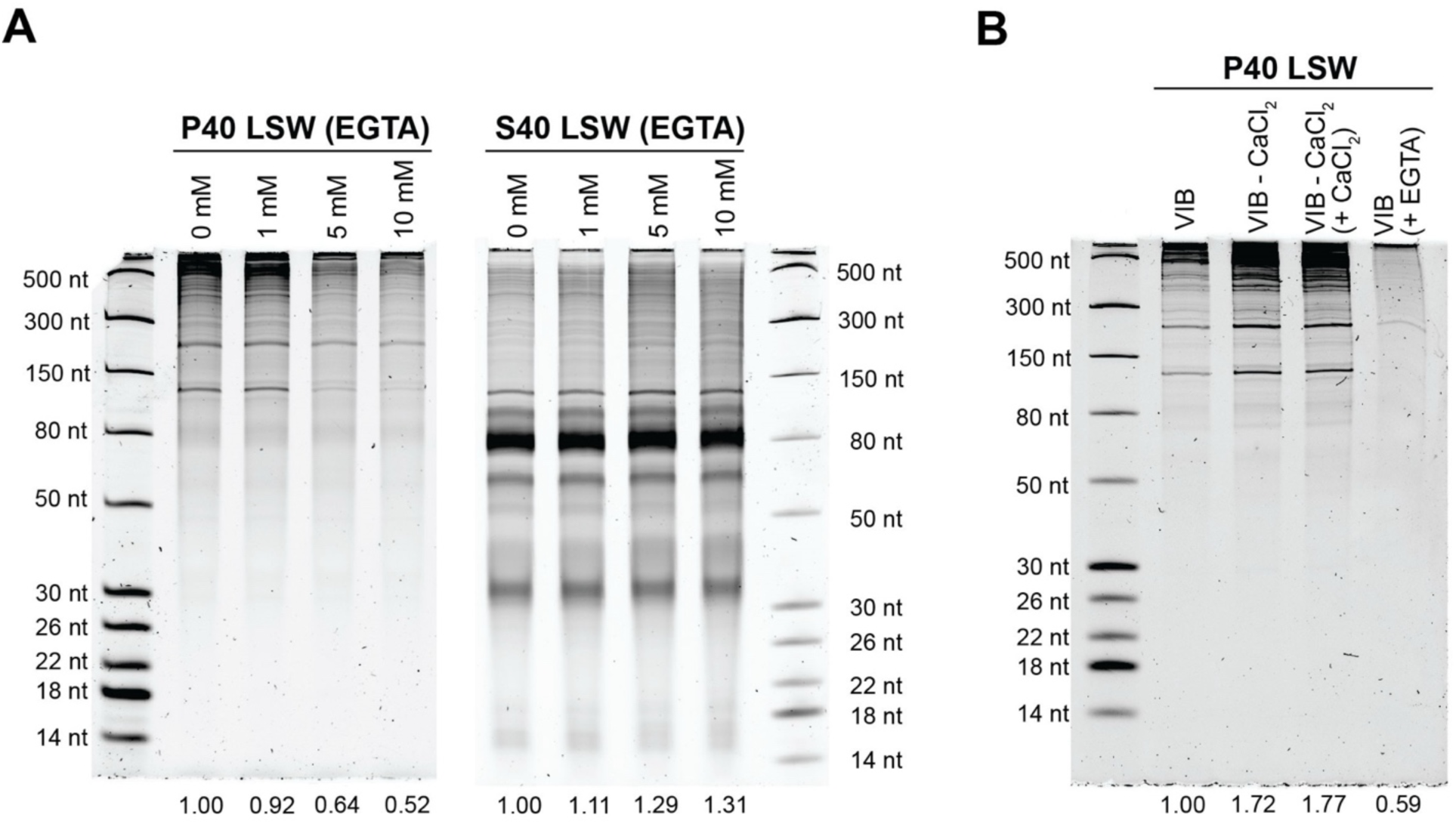
Long RNAs in LSW form cation-dependent aggregates or condensates. (*A*) An independent biological replicate of the experiment presented in Fig.7*B*. LSW was treated with increasing concentrations of EGTA as indicated and incubated on ice for 20 min, followed by ultracentrifugation at 40,000 *g*. RNAs were isolated from P40 pellets and their corresponding supernatants using TRIzol, separated on a 15% denaturing polyacrylamide gel, and stained with SYBR Gold nucleic acid stain. RNA abundance in each gel lane was estimated by densitometry using ImageJ and expressed relative to the 0 mM EGTA lane. (*B*) Gel showing the RNA profile of P40 pellets obtained from LSW fractions that were isolated using either VIB or VIB without CaCl_2_ (VIB-CaCl_2_). LSW samples were subjected to the treatments indicated in brackets for 20 min on ice prior to ultracentrifugation at 40,000 *g*. RNAs were isolated from P40 pellets using TRIzol, separated on a 15% denaturing polyacrylamide gel, and stained with SYBR Gold nucleic acid stain. RNA abundance in each gel lane was estimated by densitometry using ImageJ and expressed relative to the VIB lane.

**Fig. S14.**
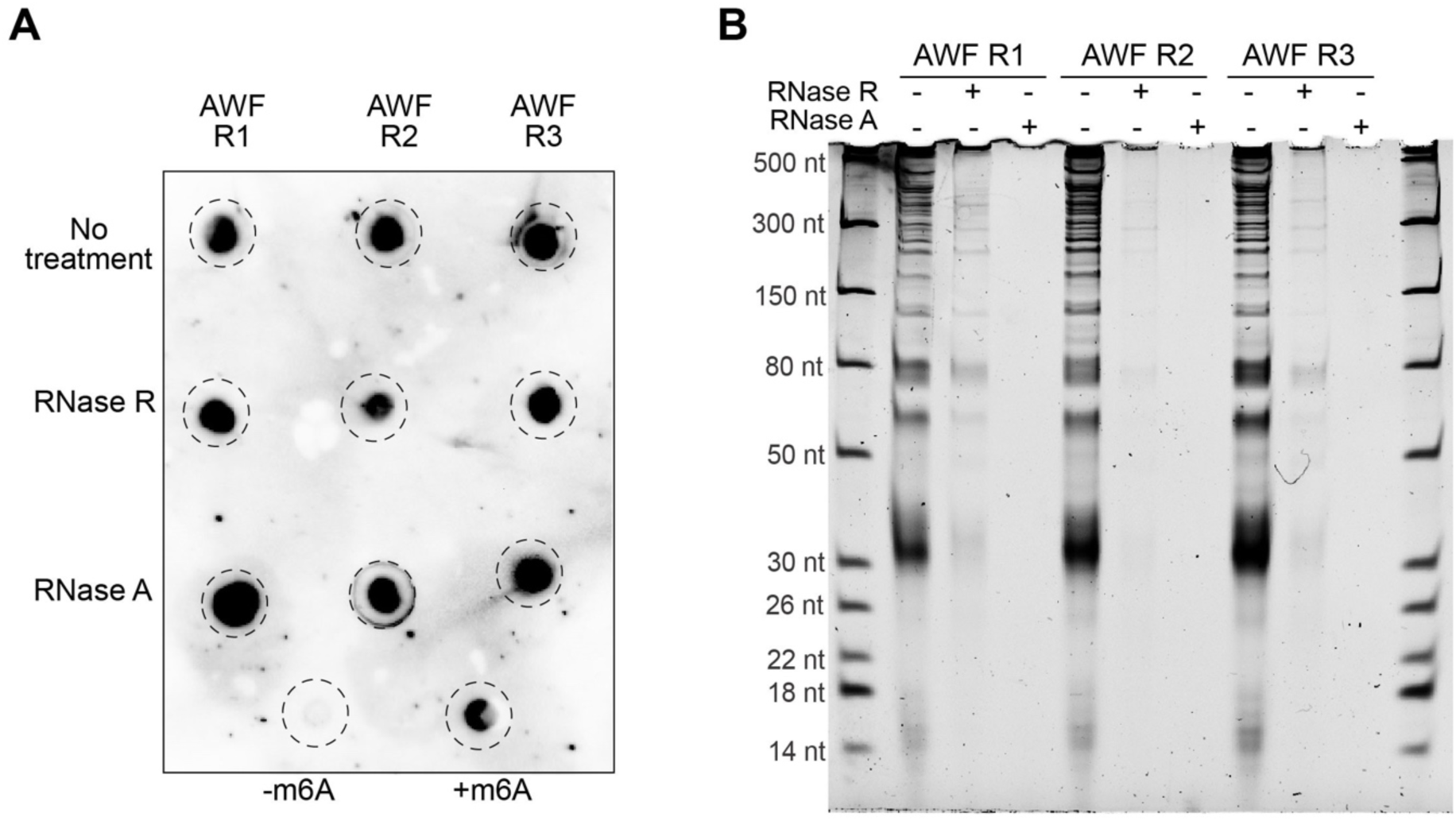
**m6A antibody cross reacts with some molecule other than RNA**. *(A)* 200 ng of AWF RNA was either not treated or treated with RNase A or RNAse R, followed by two serial precipitations with ammonium acetate and EtOH to remove free nucleotides and other small degradation products. The pellets were resuspended in 8 µL of ultrapure water, and 6 µL were dot blotted and crosslinked onto a positively charged nylon membrane and then probed with an anti-m6A antibody. Three biological replicates (R1, R2 and R3) were analyzed. For positive and negative controls, 600 ng of synthetic 21-nt RNAs with identical sequences (except for a single m6A modification on the positive control) were used. *(B)* 2 µL of RNA left from A) were separated on 15% denaturing polyacrylamide gel and stained with SYBR Gold nucleic acid stain.

**Table S1.**
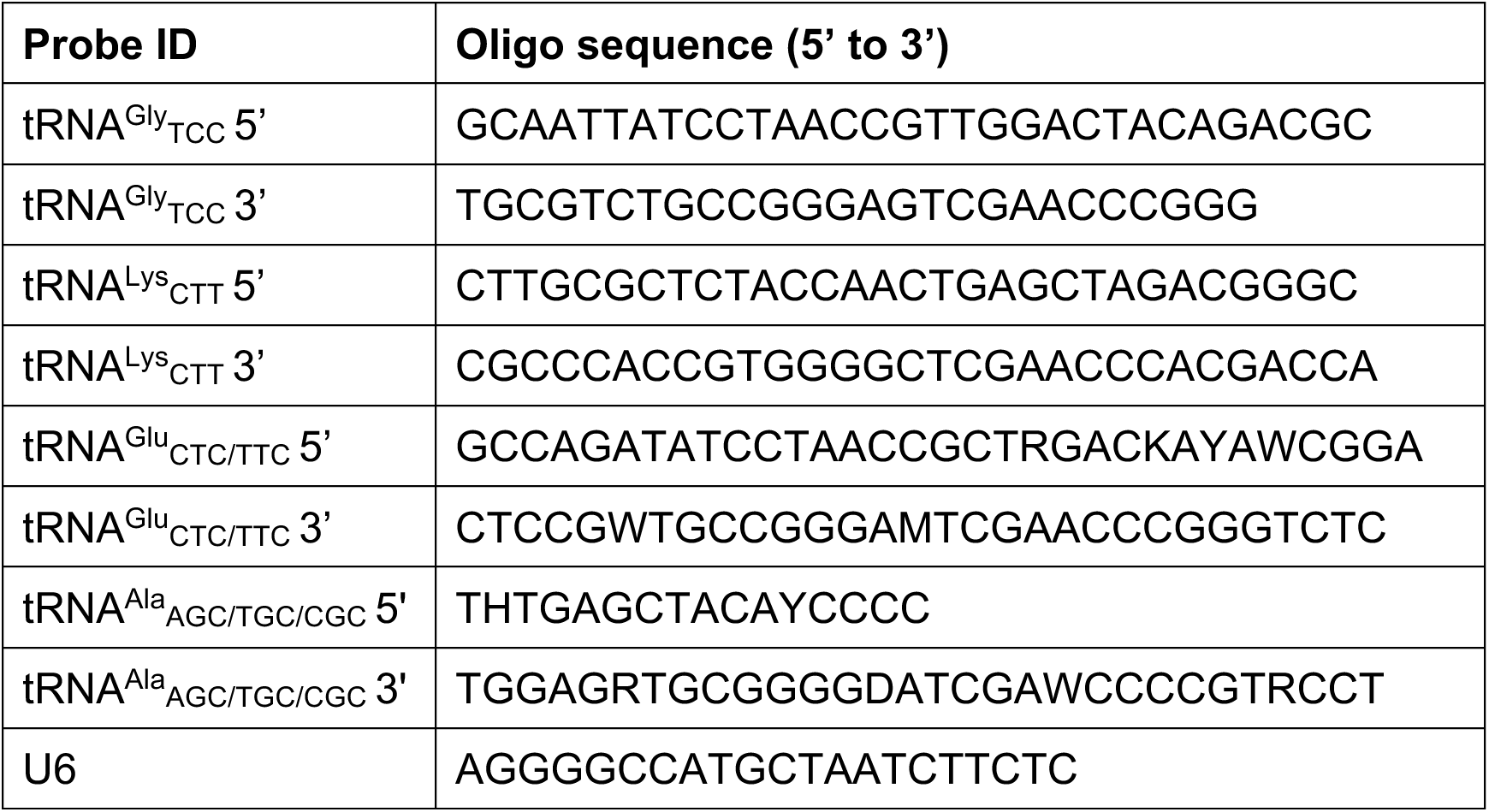
Oligonucleotide sequences of hybridization probes.

## Supplemental Datasets

**Dataset S1**. tRNA sources of tRNA-derived fragments identified in sRNAseq dataset

**Dataset S2.** Genes displaying differential transcript abundance between apoplastic wash fluid (AWF), leaf surface wash (LSW), and/or total cell lysate (CL)

**Dataset S3.** Gene Ontology analysis of differentially expressed genes in apoplastic wash fluid (AWF), leaf surface wash (LSW), and total cell lysate (CL)

